# UV induces common cutaneous amyloid-like melanosomal protein aggregates

**DOI:** 10.64898/2025.12.17.689083

**Authors:** Nicholas Theodosakis, Porter B. Howland, Miroslav Hejna, Stephen M. Ostrowski, Talya A. Cohen-Neamie, Remi A. Joseph, Helen Ji, Allison E. Greuel, Judith R. Boozer, Mariam Y. Demian, Alin Asim, Khanh Nguyen, Andrew Mu, Philip Seifert, Konrad Kaminiow, Yifan Zhang, Aline Fassini, Sharon Germana, Kristin C. Shaw, William M. Lin, Mai P. Hoang, Xunwei Wu, Sreeganga S. Chandra, Nir Hacohen, Andrea Ballabio, Rachit Bakshi, Namrata D. Udeshi, Anand K. Ganesan, Thomas Horn, George F. Murphy, Steven A. Carr, Bradley T. Hyman, Jun S. Song, David E. Fisher

## Abstract

Misfolding of aggregation-prone proteins underpins diseases known as proteinopathies. One of these proteins, alpha-synuclein, is a component of aggregates in neurodegenerative conditions such as Parkinson’s disease. The melanosomal protein PMEL, which forms physiologic amyloid scaffold structures on which melanin is organized in melanosomes, similarly ectopically accumulates in the dermis in many forms of cutaneous hyperpigmentation. Here, we demonstrate in a wide range of common clinical pigmentary disorders, as well as in primary melanocyte and mouse models examined by molecular, proteomic, and electron microscopic tools, that melanocytic alpha-synuclein is a prominent component of intracellular protein aggregates bound to similar proteins as in Parkinson’s disease, as well as melanized extracellular protein deposits. Using the Real Time Quaking-Induced Conversion Assay (RT-QuIC), we demonstrate that UV induces misfolded melanosomal proteins to self-propagate, augmenting this pathology in prion-like fashion. CUT&RUN chromatin profiling and single-cell RNA-seq demonstrate that melanocytes utilize microphthalmia-associated transcription factor (MITF)-regulated autophagy to counteract protein aggregation, identifying aggregate removal as a core function of tanning. In contrast to extracellular aggregation, impaired intracellular aggregate removal contributes to melanocyte senescence, which conversely exacerbates chronic hypopigmentation and photoaging-related discoloration. These findings identify melanosomal proteinopathy as a common contributor to melanocyte dysfunction and suggest aggregate-focused management approaches.

## Introduction

Pathophysiologic amyloid formation underpins a family of diseases of protein aggregation sometimes referred to as proteinopathies^1–3^. This includes neurodegenerative disorders such as Parkinson’s disease (PD) or Alzheimer’s disease (AD), where misfolding and insolubilization of intrinsically amyloidogenic proteins are associated with loss of function^4^. A primary component of aggregates in PD, Lewy Body Dementia, and other central nervous system (CNS) diseases is alpha-synuclein (SNCA): a 14kDa intrinsically disordered neuronal protein which possesses innate self-aggregatory potential, through which misfolded species seed misfolding and aggregation of native monomers in a prion-like fashion^5^. This process is thought to be potentiated by exposure to reactive oxygen species (ROS), as well as defects in aggregate clearance, leading to persistence of insoluble deposits that accumulate over time^6,7^.

Chronic hyperpigmentation, caused by triggers including age, sun damage, inflammation, and toxin exposure^8^, often shows a common histologic endpoint characterized by accumulation of ectopic dermal pigment deposits^8^: a state sometimes referred to as “dermal hyperpigmentation.” Some of these deposits appear to be extracellular, while others collect within skin-resident macrophages, sometimes termed “melanophages.” In addition to melanin, dermal deposits often contain multiple melanosomal proteins^9,10^, suggesting that impaired protein removal may contribute to this pathophysiology as in CNS proteinopathies. Indeed, diseases with impaired lysosome function, such as GM1 gangliosidosis and Hurler syndrome, are also known to exhibit various forms of widespread hyper– or hypopigmentation, supporting the notion that protein aggregate removal is also key to proper pigment homeostasis^11–13^. Interestingly, previous studies of sporadic PD patients have suggested that they may show both accelerated hair graying as well as decreased tanning ability^14^, promoting the concept that conditions which contribute to PD proteinopathy may also exacerbate pigmentary pathology.

As melanosomes mature, they synthesize melanin around a fibrillar protein scaffold composed of premelanosomal protein (PMEL, also known as gp100 or SILV)^15,16^. This scaffold is an example of a physiologic amyloid, which self-assembles inside melanosomes into the hallmark cross-β sheet structure, organizing melanin polymers for transfer^17,18^. Interestingly, SNCA has also recently been found to be expressed in melanocytes as a target of MITF: the master tanning transcription factor^19,20^. UV exposure activates MITF, driving transcription of these and hundreds of other genes required for pigmentation, significantly increasing the levels of these aggregation-prone proteins in hyperpigmented skin while simultaneously generating pro-aggregatory ROS. Similarly, increased SNCA levels due to gene duplication is known to underpin many forms of familial PD^21^, while age-related accumulation of amyloidogenic transthyretin or β2m in the blood contribute to cardiac and dialysis-related amyloidosis^22,23^, further supporting the hypothesis that hyperpigmentation may represent a common transient pro-amyloidogenic state.

Given these similarities of dyspigmentation to known proteinopathies, we sought to investigate whether aggregation of melanosomal components might contribute to pigmentary dysfunction. We provide evidence that oxidative stressors, including UVA radiation, greatly increase the aggregation of certain pigment proteins. This results in two separate dyspigmenting effects: triggering hyperpigmentation via the formation of insoluble pigment and melanin complexes extracellularly while intracellularly suppressing melanocytic function via senescence, leading to long-term hypopigmentation. This is also a significant public health concern given that UVA comprises more than 90% of solar UV radiation, more efficiently generates ROS than UVB, and is often only partially filtered out by commercial sunscreens^24,25^. Taken together, this study suggests that dyspigmentation may represent a form of melanosomal proteinopathy, providing a rationale for exploration of aggregate-targeted strategies for prevention and treatment, and potentially offering novel insights into the biology of proteinopathies in the CNS and other organ systems.

## Results

### Pigment proteins aggregate in darkened skin

In biopsies of hyperpigmenting disorders, we observed numerous thioflavin-positive objects in the dermis, suggesting amyloid and other protein aggregates are a common finding in hyperpigmented skin (Fig. 1a). Using immunostaining with an aggregate-specific antibody, we also observed epidermal and dermal enrichment and colocalization of aggregated SNCA with other melanosomal proteins, including PMEL and tyrosinase, confirming that many of these objects are derived from melanosomal components and that PMEL and SNCA colocalize extracellularly (Fig.1b, Extended Data Fig.1a,b). Significant amounts of aggregated SNCA also appeared to be internalized by dermal macrophages in hyperpigmented skin, which were often observed near thioflavin-positive signal (Fig. 1c, Extended Data Fig. 1c). Lack of MITF target expression changes with overexpression of SNCA in SK-Mel-30 melanoma cells supported that observed SNCA enrichment likely reflected post-translational activity (Extended Data Fig. 1d).

**Fig. 1:**
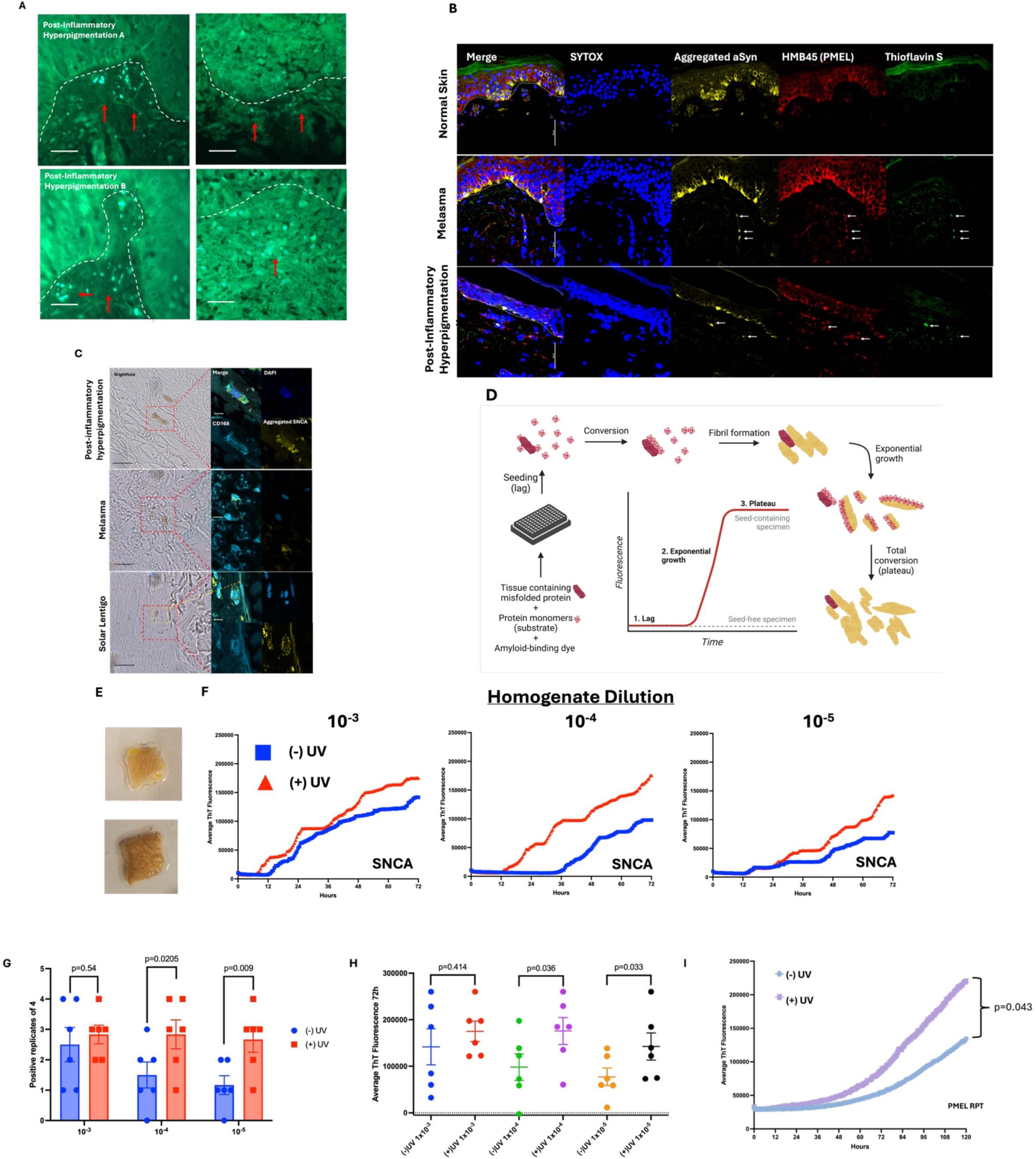
Melanosomal proteins released into the dermis spontaneously self-aggregate in hyperpigmented skin. (A) Thioflavin S staining of biopsies of various etiologies of hyperpigmented skin consistently shows positive objects in the superficial dermis (red arrows), dermal-epidermal junction (DEJ) is labeled with a white dashed line, scale bar = 20µm. (B) Representative images of thioflavin and immune co-staining of patient biopsies show that dermal amyloid objects also stain positively for SNCA using an aggregate-specific antibody and for PMEL (arrows), scale bar = 50µm. (C) Aggregated SNCA (yellow) is often found within and in close proximity to dermal macrophages (CD168, light blue) (red boxes), as well as extracellular melanin (orange box) in hyperpigmented skin (black bar = 10µm, white bar = 2µm). (D) The Real-Time Quaking-Induced Conversion Assay measures the ability of protein in tissue homogenate to induce misfolding and aggregation of recombinant soluble monomers in prion-like fashion, which is translated into an optical signal by thioflavin dye. (E) Discarded skin from routine abdominoplasties from a total of 6 donors was divided and irradiated with 5J/cm^2^ UVA (peak em. 368nm) daily for 10 days. UVA tanning is known to primarily be driven by melanin oxidation rather than tanning pathway induction. (F) Serial dilution of tissue homogenate was assay by RT-QuIC using recombinant SNCA as substrate. Lines represent an average fluorescence value across four technical replicates from all six donors at 30 minute intervals across 72 hours. (G) Average fluorescence for all technical replicates across all 6 donors at the 72 hour endpoint are depicted. Two-tailed paired student’s *t-*test was used to compare unirradiated vs irradiated skin at each dilution. (H) The number of positive wells at 72 hours was tallied for each donor at 72 hour endpoint. Positive wells were defined as those whose signal exceeded 10% of maximum signal at least twice. Positive well counts for unirradiated vs. irradiated skin were compared by Pearson’s ***x***^2^ test. (I) RT-QuIC was also performed on irradiated vs. unirradiated skin homogenate diluted by a factor of 1×10^−5^ from three separate donors using recombinant PMEL RPT domain as substrate. The difference in endpoint fluorescence is depicted at the 120 hour assay endpoint using two-tailed paired student’s *t-*test.

A hallmark of CNS synucleinopathies is thought to be propagative formation of protein aggregates over time in a manner sometimes likened to prion disease^5,26^. We asked whether skin-endogenous SNCA may show similar activity. Using the highly sensitive Real-Time Quaking-Induced Conversion Assay (RT-QuIC)^27–29^ (Fig. 1d), we UVA-irradiated human skin explants from discarded surgical specimens for 10 days and measured self-aggregatory activity (Fig. 1e, Extended Data Fig. 2). Given known links between oxidative stress in the brain and protein misfolding^30^, UVA was chosen due to its ability to efficiently generate ROS without significantly damaging DNA or crosslinking protein compared to UVB^31^. UVA is also known to minimally activate MITF-induced tanning, though it does cause skin darkening due to oxidation of pre-exiting melanin^32^. UV-treated skin consistently showed significantly greater activity as compared to non-irradiated skin (Fig. 1f-h, Extended Data Fig. 2). A similar pattern was observed with RT-QuIC performed using recombinant PMEL RPT domain as substrate, which has also been reported to show self-aggregation (Fig. 1i)^17,33,34^.

To further explore the physiologic importance of synuclein family members to pigmentation, we irradiated germline *Snca*^KO^ and alpha-, beta-, and gamma-synuclein triple knockout mice (*Snca/b/g*^KO^) to induce tanning^35,36^. At baseline, these mice lack significant coat or ear color defects (Fig. 2a-d) but show 27% less ear darkening with daily UVB exposure compared to WT controls (Fig. 2d,e). *Snca*^KO^ and *Snca/b/g*^KO^ mice also failed to show any significant increase in dermal melanophages with UVB irradiation (Fig. 2f), supporting that SNCA contributes at least in part to dermal pigment accumulation.

**Fig. 2:**
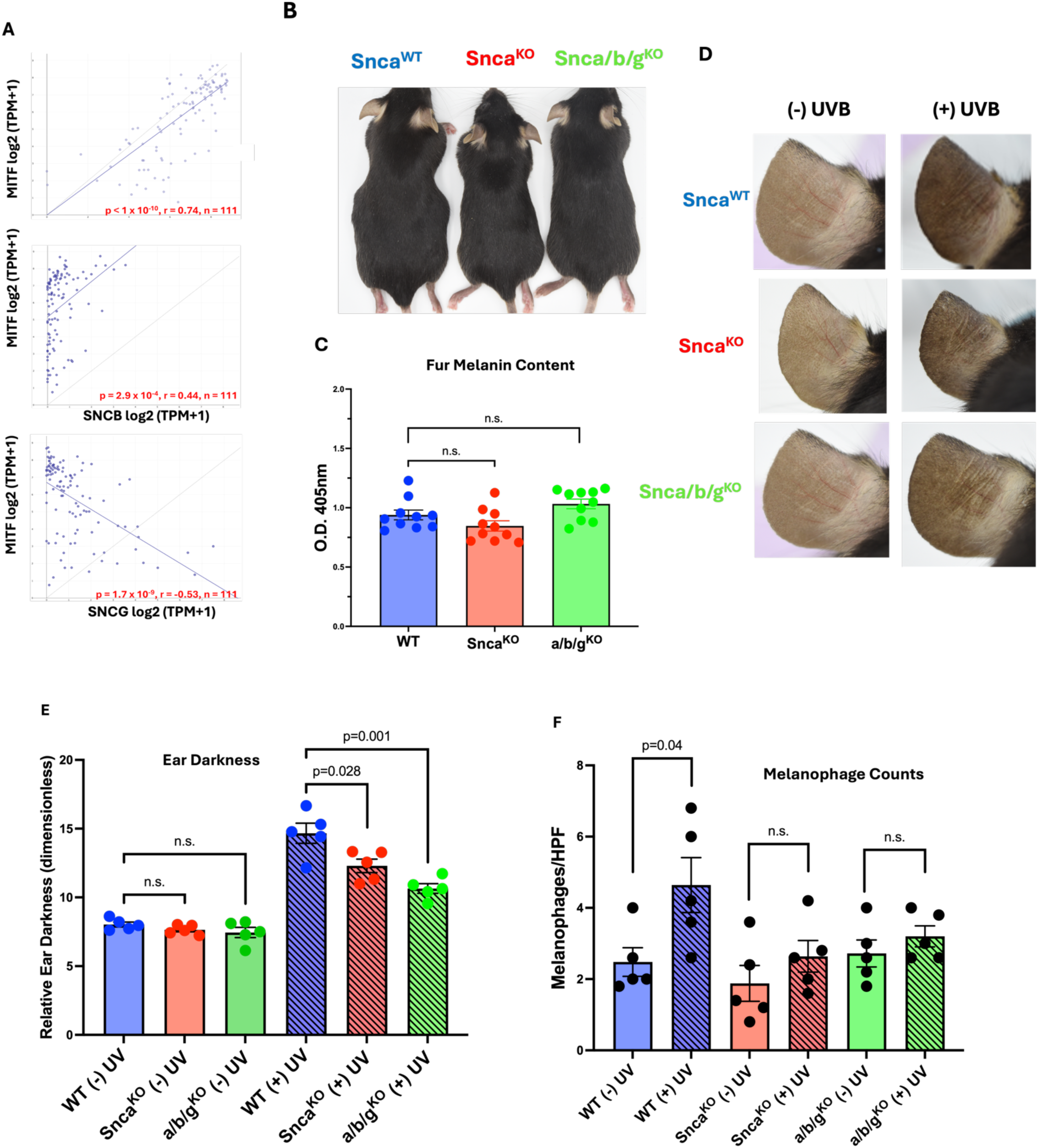
Loss of synuclein family members impairs UV tanning and melanophage formation. (A) Expression of MITF and synuclein family members in melanoma cell lines in the Cancer Cell Line Encyclopedia database shows a positive correlation between levels of MITF and SNCA and SNCB but a negative correlation with SNCG (CCLE Expression Public 25Q3). (B) C57BL/6 mice with germline deletion of *Snca* or deletion of alpha-, beta-, and gamma-synuclein do not show defects in coat or ear coloration at baseline. (C) Single and triple knockout mice show no significant difference in fur melanin content (student’s *t*-test, n=10). (D,E) Exposure to 100mJ/cm^2^ UVB radiation (peak em. 302nm) 5 days/week for 4 weeks resulted in decreased ear darkening in knockout mice vs. WT (student’s *t-*test, n=5). UVB was used rather than UVA due to the minimal tanning response generated by UVA radiation. (F) Melanophage formation above baseline with UVB exposure was also decreased in knockout mice vs. WT controls (student’s *t*-test, n=5).

### ROS exacerbates protein aggregation

Self-propagated protein aggregation in PD, as measured by RT-QuIC, is often modeled using SNCA overexpression mouse strains^37^. To model the melanosomal protein aggregation suggested by our thioflavin staining and skin RT-QuIC data *in vivo*, we generated a mouse strain targeting PD-like SNCA overexpression within skin melanocytes. This included the K14-SCF mouse transgene, which confers human-like interfollicular epidermal melanocytes, and a melanocyte-specific tetracycline-inducible promoter linked to dopachrome tautomerase^38,39^. These were combined with overexpression constructs for WT human SNCA (*K14-SCF:Dct-rtTA:SNCA^WT^*, termed “SNCA^WT^” mice) or SNCA harboring an A53T mutation (*K14-SCF:Dct-rtTA:SNCA^A53T^* termed “SNCA^A53T^” mice), which resulted in overall SNCA levels approximately 3-fold higher than basal Snca expression (Methods). The A53T mutation significantly increases SNCA’s aggregatory potential and underpins many kindreds of familial Parkinsonism^39,40^. Importantly, WT mouse Snca has a threonine at this residue, increasing human SNCA^A53T^’s ability to induce misfolding in endogenous mouse synuclein despite the known species barrier^41^. Given the established contribution of oxidative stress to CNS SNCA aggregation, we treated mice with either topical 10% benzoyl peroxide (BPO) or UVA irradiation, which was again chose to maximize ROS generation while minimizing tanning pathway activation, DNA mutation, or direct protein crosslinking. K14-SCF mice are also intrinsically pigmented, eliminating the need for UVB tanning induction required with non K14-SCF knockout mice.

Strikingly, doxycycline induction of WT or mutant SNCA overexpression led to the appearance of large numbers of hyperpigmented macules interspersed with patches of hypopigmentation (Fig. 3a). Hyperpigmented lesion count, though not hypopigmentation, was greatly potentiated by treatment with either BPO or UVA, with the A53T aggregation-prone mutant developing more spots and more hypopigmentation than WT with no change in epidermal melanocyte counts (Fig. 3b-f). Both models show significant enrichment of SNCA, melanin deposits, and macrophages in the dermis (Extended Data Fig. 3a-c). Given the general phenotypic similarity of SNCA^WT^ and SNCA^A53T^ and the reduced species barrier conferred by the A53T mutation^42^, we utilized the SNCA^A53T^ model to define the mechanism underlying the pigment abnormalities. Interestingly, spots were observed to be labile, fading with doxycycline and BPO withdrawal, and re-darkening at the same locations with reintroduction of doxycycline only (Fig. 3g, Extended Data Fig. 3d). Dermal macrophage counts mirrored changes in pigmentation, with macrophages increasing in hyperpigmented skin overexpressing SNCA and fading after doxycycline withdrawal (Fig. 3h). Transmission electron microscopy (TEM) of dermal melanophages in SNCA^A53T^ mice revealed phagosomes that accumulated both melanosome breakdown products and hypodense material that was absent from melanophages in control animals. This material likely represented aggregated SNCA that was detected in melanophages of SNCA^A53T^ mice by immunofluorescence microscopy (Extended Data Fig. 3e,f). SNCA^A53T^ (though not SNCA^WT^) mice also rapidly depigment new fur growth within 2 weeks of doxycycline induction, suggesting a similar effect on hair bulb melanocytes (Extended Data Fig. 3g). We conclude that SNCA overexpression causes aggregation that impairs melanogenesis in epidermal melanocytes, leading to hypopigmentation, but leads to the accumulation of extracellular melanized aggregates in the dermis and within dermal melanophages, causing focal hyperpigmentation.

**Fig. 3:**
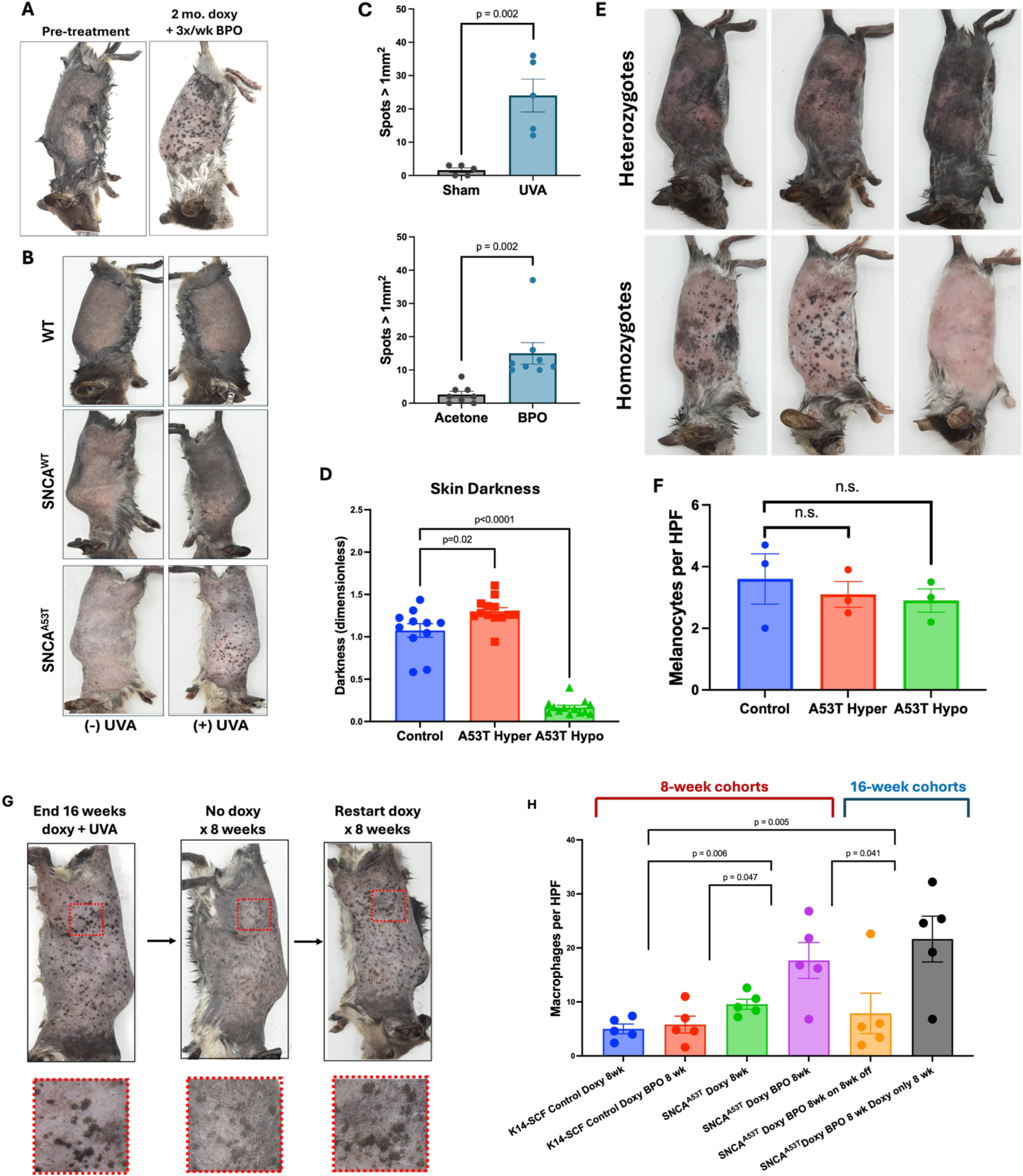
Aggregation of melanocytic alpha synuclein induces pigmentary dysfunction and melanophage formation. (A) Induction of synuclein overexpression triggers pigmentary dysfunction which is exacerbated by oxidative stress. (B) exposure of unilateral flank skin to 5J/cm^2^ 3x/week UVA irradiation (peak 368nm) for 16 weeks failed to tan WT mice significantly but greatly potentiates hyperpigmented spot development vs. the UV-protected opposite flank. (C) Mouse flanks exposed to oxidative stress in the form of UVA irradiation (5J/cm2, 3x/week) or 10% w/v benzoyl peroxide develop larger numbers of spots than UV-protected or acetone vehicle-treated flanks in split flank studies (n≥5 mice, two-tailed student’s *t*-test, p<0.01). (D) Dark spots on SNCA^A53T^ mice were modestly darker than mice fed control chow, but hypopigmented areas showed a profound decrease in pigment levels as measured by photography (n = 11, student’s *t*-test). (E) SNCA^A53T^ mice homozygous for the Dct-rtTA and SNCA^A53T^ transgenes showed more pronounced hyper– and hypopigmentation vs. heterozygotes after 8 weeks of 3x weekly BPO treatment and doxycycline induction. (F) Sox10 staining of SNCA^A53T^ skin showing no significant change in epidermal melanocyte count in hyper– or hypopigmented skin (two-tailed student’s t-test, n=3 mice). (G) After spots develop in response to doxycycline induction and oxidative stress, spots gradually fade with cessation of doxycycline and re-darken with reintroduction of doxycycline without additional oxidative stress. (H) Administration of doxycycline chow with 3x-weekly topical BPO vs acetone vehicle yielded an increase in dermal macrophages which reversed with doxycycline and BPO withdrawal (n=5 mice per condition, student’s two-tailed *t*-test)

To test whether oxidative stress enhances the intracellular accumulation of misfolded protein within melanocytic cells, we treated SK-Mel-30 melanoma cells in culture with rotenone (to induce ROS) and chloroquine (to inhibit autophagy) as has previously been described in the PD literature^43,44^. Rotenone treatment alone led to a significant increase in per-cell misfolded protein burden as measured by the proteostat assay which further increased with chloroquine^45^ (Fig. 4a). Similarly, rotenone/chloroquine treatment increased the accumulation of SNCA in triton-insoluble fractions of SK-Mel-30 cells (Fig. 4b). This includes a greater proportion of SNCA monomer as well as higher MW bands and smear, both supporting that a combination of ROS and autophagy inhibition lead to the accumulation of SDS-resistant SNCA multimers or complexes with other proteins. The accumulation of a 100 kDa fragment in the detergent-insoluble fraction of rotenone-treated SK-Mel-30 cells also suggests that ROS impairs PMEL proteolysis, a critical requirement for PMEL fibril formation, which might contribute to extracellular pathologic PMEL aggregation^16,46,47^. This interpretation was also supported by an increase in SNCA in intermediate density sucrose fractions of primary melanocyte lysate after reversible DSP crosslinking, also suggesting separation of higher MW aggregates prior to cleavage (Fig. 4c). To test whether similar synuclein aggregation was induced by UV radiation, we irradiated primary human melanocytes with 5J/cm^2^ daily UVA (approximately equal to 10 minutes of midday sun on a clear day in the United Kingdom^48^) for 3 days. This led to formation of small aggregated SNCA-positive inclusions (Fig. 4d,e). Co-staining of aggregates with p62 suggests that many of these aggregates may be enclosed within autophagosomes, also suggesting attempted clearance (Fig. 4f).

**Fig. 4:**
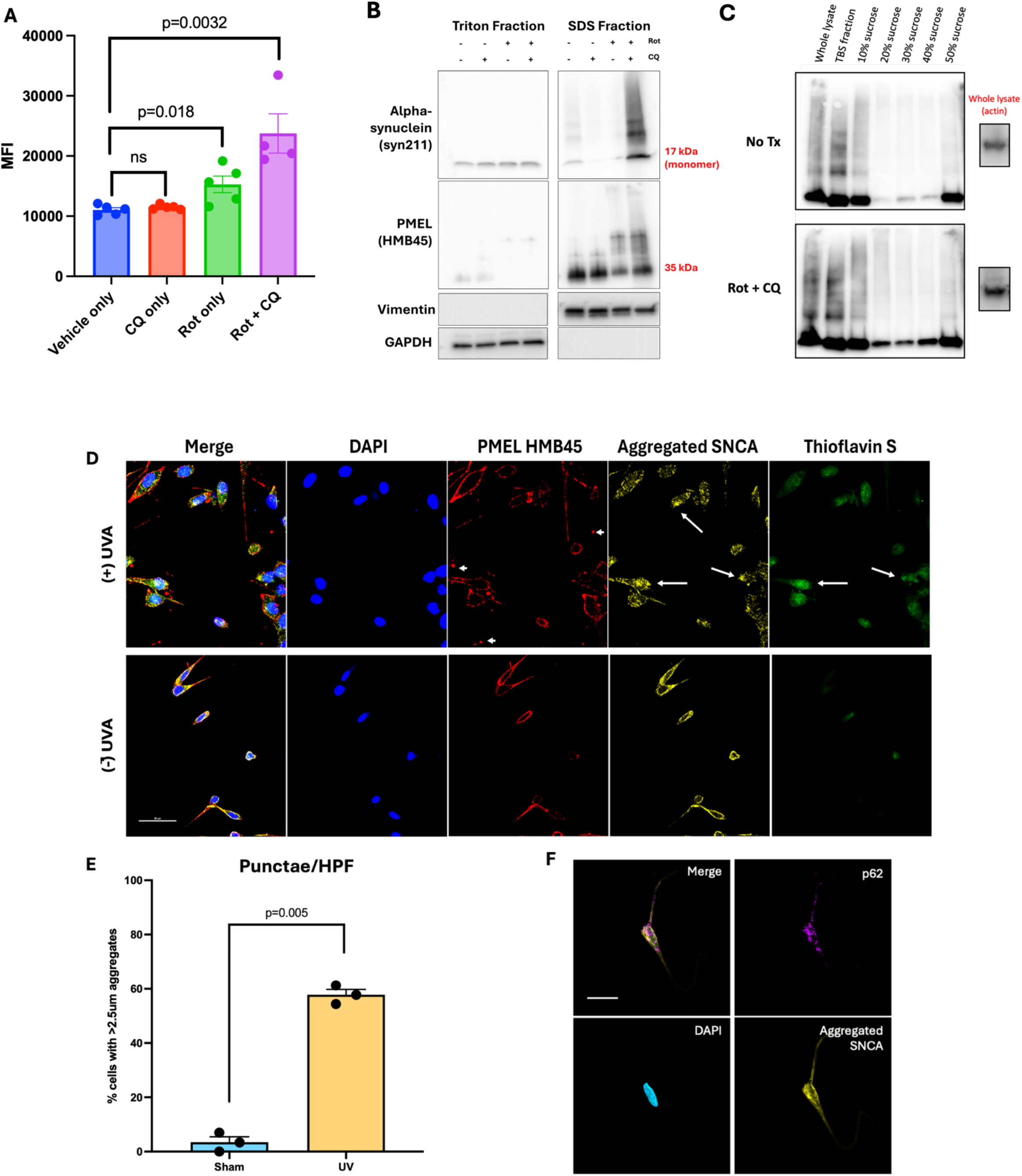
ROS induces intracellular SNCA aggregation. (A) Treatment of SK-Mel-30 melanoma cells with 100nm rotenone for 72 hours and 20µm chloroquine for 24 hours induced a significant increase in total misfolded protein as assayed by flow cytometry with staining using the proteostat aggregation dye (n=5, student’s *t*-test). (B) ROS induction through treatment with 100nm rotenone for 72 hours with co-treatment with 24h 20µM chloroquine leads to significant insolubilization of both PMEL and SNCA and enrichment of higher MW forms. (C) Identical treatment of primary melanocytes with rotenone and CQ followed by protein crosslinking with the cleavable crosslinker DSP and ultracentrifugation through a discontinuous sucrose gradient showed enrichment of SNCA in intermediate fractions in treated samples vs. control before cleavage with BME. (D) daily 5J/cm^2^ UVA irradiation of human primary melanocytes for three days leads to the formation of SNCA-positive inclusions (arrows), some of which also stain positive for amyloid. Extracellular PMEL-positive objects are also seen (arrowheads), though minimal intracellular colocalization of SNCA and PMEL was observed (E) Significantly more SNCA-positive punctae are seen in these UV-irradiated cells compared to sham-irradiated melanocytes (n=3 experiments, average of 5 fields per experiment), student’s *t*-test). (F) UV-irradiated primary melanocytes show colocalization of SNCA and p62, suggesting autophagy-mediated degradation of damaged proteins

To study the relevance of SNCA prion-like activity in vitro, we generated amyloidogenic pre-formed SNCA fibrils (PFFs) as previously described^49^ and transfected them into primary human melanocytes. Co-treatment with bafilomycin to inhibit autophagy resulted in progressive formation of high MW insoluble SNCA species (Extended Data Fig. 4a). Notably, PFF-induced SNCA aggregation did not induce a significant increase in insoluble PMEL, supporting our previous ICC data suggesting that the two species do not directly interact intracellularly. RT-QuIC analysis of lysate from UVA– and bafilomycin-treated primary melanocytes showed significant positivity in the UV+bafilomycin condition (Extended Data Fig. 4b,c), supporting a role for spontaneous aggregation following UV exposure as well as an active role for autophagy in counteracting this UV-induced process. We also observed that solubilizing primary melanocyte pellets in SDS released increasing amounts of melanin into supernatant lysate (Extended Data Fig. 4d-f), suggesting that a significant fraction of melanin may be bound to insoluble protein in pathologic hyperpigmentation^15,50^.

To identify proteins that bind to SNCA under pathological conditions, we used a proteomics analysis of SNCA immunoprecipitates. SK-Mel-30 melanoma cells that overexpressed SNCA were treated with rotenone and with bafilomycin A1, which inhibits autophagy by impairing lysosomal acidification^51,52^. SNCA was immunoprecipitated from these cells, and co-immunoprecipitated proteins were analyzed by tandem mass spectrometry compared to CRISPR *SNCA*^KO^ controls. The results showed significant enrichment for proteins involved in membrane formation and transmembrane transport, including known pigmentation pathway targets (Extended Data Fig. 5a-f, Supplementary Table 1). This mirrored a previous mass spectrometry study of SNCA interactors in PD neurons that also identified proteins involved in vesicle trafficking and transmembrane protein transport^53^.

### MITF regulates tanning byproduct removal

In melanocytes, autophagy has been demonstrated to be involved in processes related to pigmentation, including melanosome trafficking and delivery of synthetic enzymes^54–56^. Less attention has been given to autophagy’s role in elimination of pigmentation byproducts. This is of particular interest given that MITF-related transcription factors TFEB and TFE3 are known to regulate aggregate removal in most non-melanocytic cell lineages through activation of the Coordinated Lysosomal Expression and Regulation (CLEAR) network^57,58^. MITF has also been shown to bind to promoters of genes in the CLEAR network and to induce expression of autophagy-related genes, the functional significance of which has primarily been studied in the context of starvation adaptation using melanoma or non-melanocytic cell lineages^58–60^.

To study the importance of autophagy in SNCA aggregate clearance, we used a split luciferase SNCA expression construct^61^ and genetic ablation of autophagy with CRISPR-Cas9 deletion of ATG7 in UACC257 melanoma cell to show diminished ability to eliminate aggregates after 24 hour rotenone treatment (Fig. 5a,b). We then bred *SNCA^A53T^* mice with an *Atg7*^flox/flox^ mouse strain together with melanocyte-targeted Cre recombinase linked to the tyrosinase promoter^62^. Disruption of autophagy in vivo resulted in grossly normal pigmentation at baseline, but more severe hypopigmentation upon doxycycline induction and BPO treatment, with rare, patchy dark zones as opposed to the well-defined spots observed in Atg7^WT^ mice (Fig. 5c). Histology of dark patches revealed enlarged pigment granules in melanocytes in the epidermis which were not present in uninduced mice, as well as a paucity of dermal pigmentation (Fig. 5d), further supporting that dark spots in SNCA^A53T^ mice are comprised in large part of dermal pigment deposits. These findings also support the conclusion that when autophagy is functional, large intracellular protein aggregates are effectively broken down or exported, explaining why large pigment aggregates in hyperpigmented skin are primarily extracellular and dermal as opposed to intracellular within melanocytes. TEM of flank skin from SNCA^A53T^:Atg7^flox/flox^ mice showed basolateral localization of melanosomes, also suggesting a possible export defect induced by SNCA overexpression and oxidative stress with defective autophagy (Fig. 5e).

**Fig. 5:**
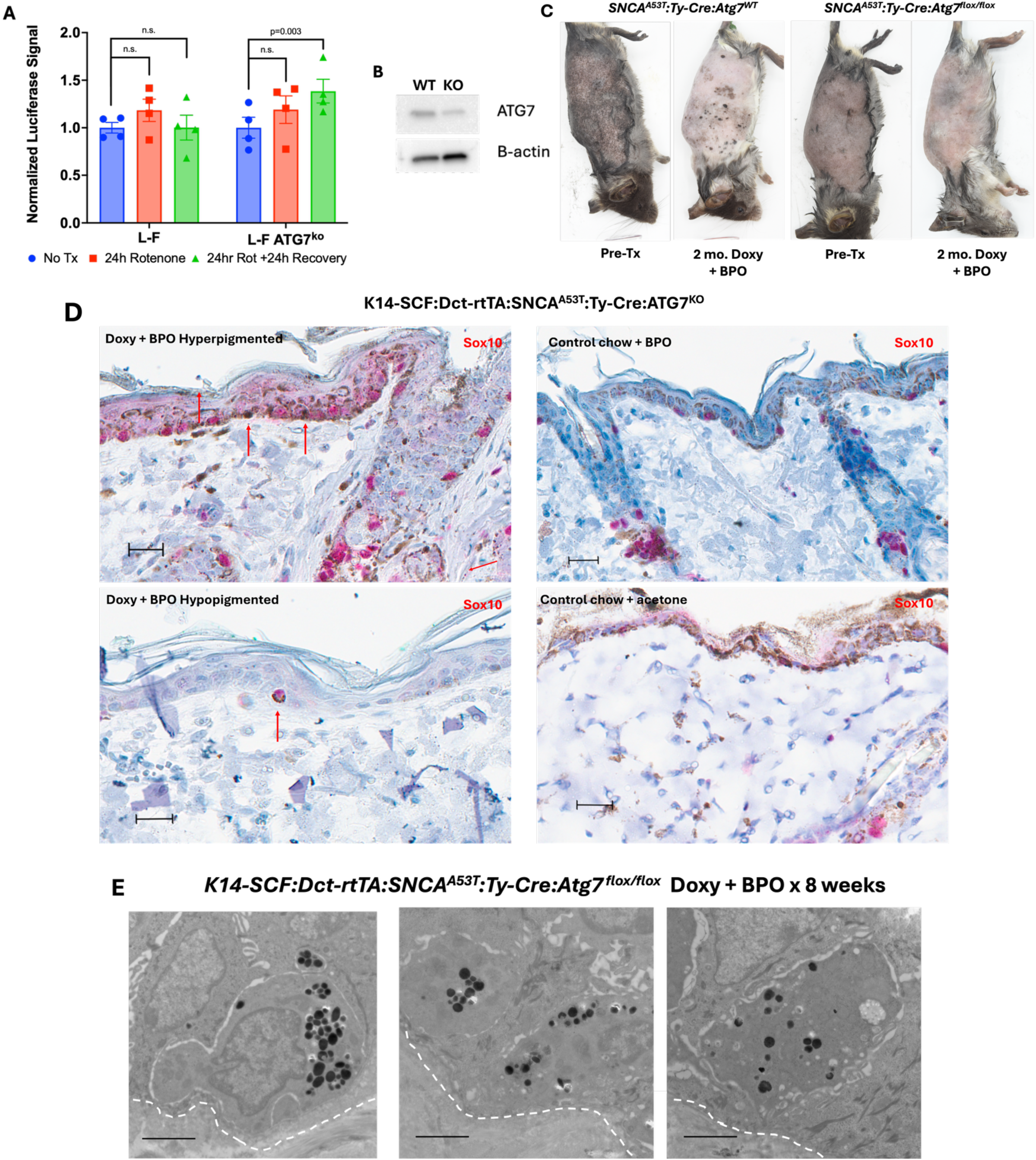
Autophagy is needed for elimination of protein aggregates and pigment homeostasis. (A,B) UACC257 melanoma cells were transduced with the split luciferase reporter construct and a construct expressing CRISPR/Cas9 and gRNA targeting ATG7 or nontargeting control to ablate autophagy. Both were treated with 100nM rotenone for 24 hours followed by a 24 hour washout period. The ATG7-null population showed continued increase in SNCA aggregation after removal of rotenone while autophagy-competent cells remained at baseline (n=3, students *t*-test). (C) *K14-SCF:Dct-rtTA:SNCA^A53T^:Ty-Cre:Atg7 ^flox/flox^* mice were induced with doxycycline chow for 8 weeks and had 10% BPO applied to their left flanks 3x/week. Mice showed primarily hypopigmentation with minimal spot formation compared to *K14-SCF:Dct-rtTA:SNCA^A53T^:Atg7^WT^*controls. (D) Sox10 staining showed accumulation of large melanized granules within and around melanocytes (red arrows) with overall decreased extramelanocytic epidermal melanin and decreased dermal melanin with the combination of SNCA overexpression and BPO (though not at baseline), suggesting an export defect and supporting that dark spots in Atg7^WT^ mice are due in large part to dermal melanin deposits. Atg7^KO^ mice fed control chow show grossly normal epidermal pigmentation with and without BPO. Scale bar = 20µm (E) Electron micrographs of skin from a *K14-SCF:Dct-rtTA:SNCA^A53T^:Ty-Cre:Atg7 ^flox/flox^* mouse showing unusual basolateral accumulations of melanosomes within melanocytes (black scale bar = 2µm, white dashed line = DEJ)

Hypothesizing that MITF may upregulate autophagy with tanning to eliminate harmful byproducts, we examined MiT/TFE family member essentiality in regulating autophagy genes in human primary melanocytes. CUT&RUN chromatin profiling^63–65^ and bulk RNA-seq of siRNA-treated primary melanocytes indicated that the majority of autophagy genes are regulated by MITF, with fewer regulated by TFE3 and almost none regulated by TFEB, giving evidence that MITF may largely supplant the role of these other family members in this lineage (Extended Data Fig. 6a-f). Melanocytes treated with siRNA demonstrated significant downregulation of autophagocytic flux with siMITF while activation with forskolin led to increased flux and clearance of misfolded protein after UV (Extended Data Fig. 6g-i). Similarly, BPO treatment of SNCA^A53T^ mice with melanocyte-specific Tfeb^flox/flox^ knockout^66^ failed to show phenotype modulation relative to Tfeb^WT^ (Extended Data Fig. 6j).

### Chronic proteinopathy induces senescence

We performed single-cell RNA sequencing on epidermis from BPO-treated primarily hyper– vs hypopigmented 8 month old SNCA^A53T^ mice and a K14-SCF control, at which point phenotypes were well-established (Extended Data Fig.7a-g, Supplementary Table 2). Gene ontology (GO) analysis showed differential regulation of pathways related to protein degradation, as well as genes related to neurodegenerative proteinopathies in both hyperpigmented and hypopigmented samples relative to control (Extended Data Fig.7h Supplementary Figure 1). Analysis showed comparatively few significant differences between hyperpigmented and hypopigmented samples (Extended Data Fig. 8), including in pigmentation genes, providing further evidence that hyperpigmentation is primarily driven by post-transcriptional dermal accumulation. Hypopigmentation of the overlying epidermis may also explain why dark spots are only marginally darker than normal skin of control mice despite greatly enriched dermal melanin. Interestingly genes related to UPR and autophagy in particular showed greater differential regulation in the hyperpigmented sample vs. hypopigmented. Given the approximately equal SNCA^A53T^ expression in both samples, we interpret this as suggesting that hyperpigmented melanocytes are able to maintain some baseline pigment pathway activation, resulting in persistent proteinopathic stress, whereas hypopigmented melanocytes have entered a fully senescent state, evidenced by upregulation of markers such as cdkn2a, in which pigmentation has been suspended to limit further proteotoxicity. Accordingly, we noted significant downregulation of effectors of the Integrated Stress Response (ISR), such as DDIT3 (CHOP) and EIF2AK4 (GCN) in the hyperpigmented sample which appear to normalize in the hypopigmented sample. ISR plays a well-established role in managing chronic proteinopathy through downregulation of global protein synthesis and senescence induction, with chronic activation known to contribute to the pathophysiology of neurodegenerative diseases^67–69^.

This same combination of decreased pigment and elevated senescence markers has been reported in a common human hypopigmenting lesion of unknown etiology noted nearly ubiquitously on sun-damaged skin, generally in the 5^th^ decade and beyond: Idiopathic guttate hypomelanosis (IGH)^70,71^. Histologic studies of these lesions demonstrate broadly normal numbers of melanocytes with decreased melanin^72,73^. We used scRNA-seq to compare lesional vs. perilesional expression features from four human participants with extensive IGH (Extended Data Fig. 9a-e, Supplementary Figures 2,3). GO analysis showed differential expression of genes involved in ubiquitin ligation and isopeptide bond formation, as well as cell-cell adhesion and cell junction formation. This includes SORBS2, which is known to play a role in dendrite formation and melanosome transfer^74^, supporting the model that chronic UV-induced proteinopathic stress, which may be exacerbated by buildup of melanosomal proteins through impaired export, contributes to IGH formation. Accordingly, we also saw significant upregulation of DDIT3/CHOP across all participants, further suggesting proteinopathy as a cause for hypopigmentation in IGH much like our hypopigmented mice. This upregulation vs. downregulation of CHOP in the mice, as well as the comparatively milder upregulation of CDKN2A in IGH, may suggest that IGH melanocytes are able to more successfully manage proteotoxicity in the setting of physiologic SNCA levels compared to the supraphysiologic misfolded protein load induced by overexpression. Several notable neurodegeneration-related genes were also significantly differentially regulated (Extended Data Fig. 9f).

## Discussion

These data provide a framework for how the innate pro-aggregatory potential of melanosomal proteins can result in common pathologic aggregate formation with oxidative stressors such as chronic UV exposure (Fig. 6). When misfolded protein and melanin are mislocalized to the dermal milieu, further aggregation is no longer restricted by intracellular regulatory mechanisms such as autophagy, allowing for aggregation of susceptible proteins such as PMEL and SNCA which do not interact intracellularly. In this framework, these insoluble extracellular melanin and protein deposits resist clearance by skin-resident macrophages, contributing to chronic dermal hyperpigmentation in the context of a range of disorders. This model of pigmentary disease as a form of “melanosomal proteinopathy” offers an explanation as to how post-translational pigment protein biochemistry may contribute to both pigmentary pathology and melanocyte photoaging.

**Fig. 6:**
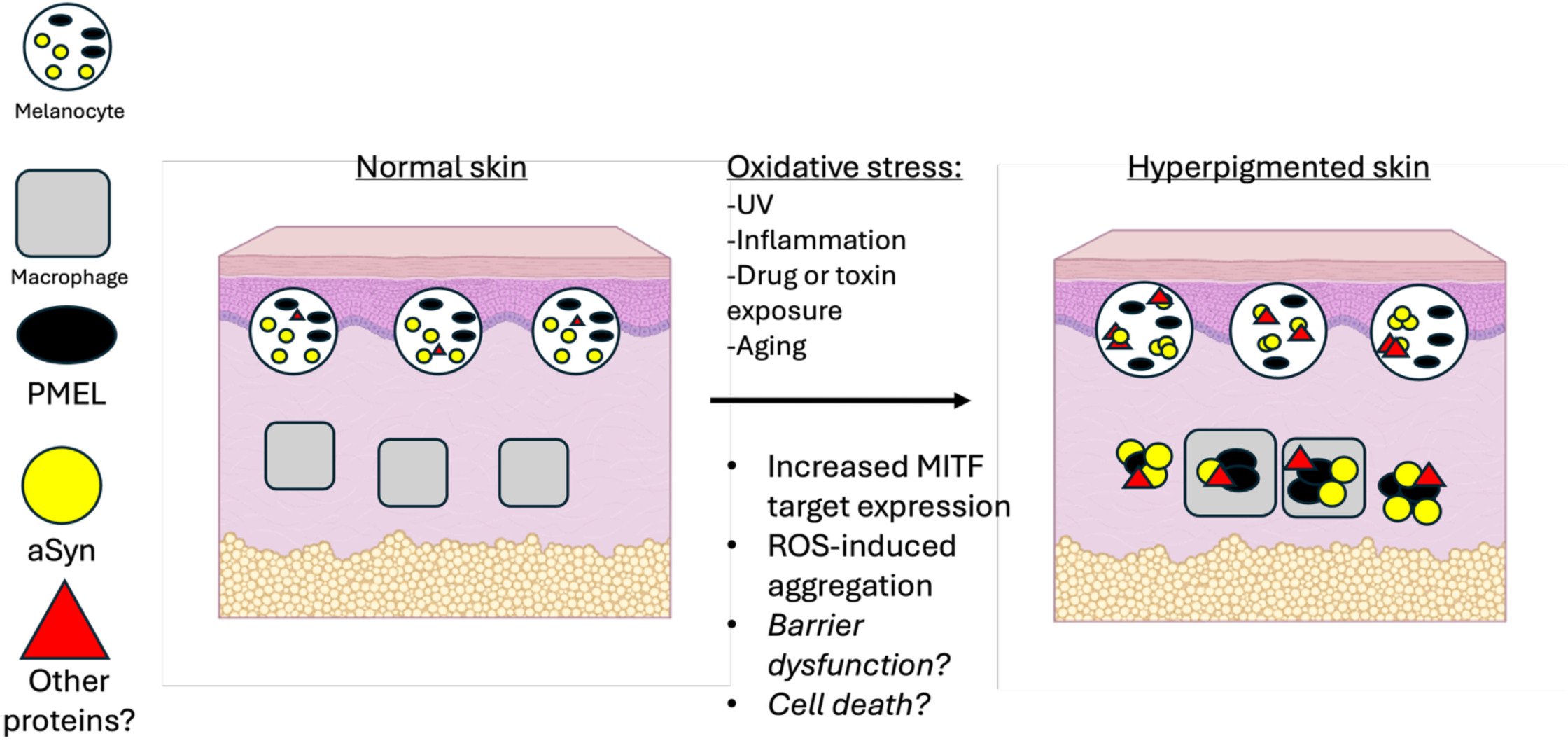
Melanosomal proteinopathy as a framework for chronic hyperpigmentation. In this model, extracellular aggregation and insolubilization of melanosomal proteins contributes to the formation of insoluble dermal deposits of protein and melanin while intracellular aggregation may sometimes contribute to melanocyte senescence and hypopigmentation from chronic proteotoxicity

This framework may also explain why chronic UV can sometimes precipitate hypopigmenting conditions such as IGH by inducing chronic intracellular as opposed to extracellular aggregation. The mouse and human scRNAseq data suggest that chronic protein misfolding triggers neurodegeneration-like changes and senescence in melanocytes, marked by UPR/ISR induction, downregulation of the tanning pathway, and upregulation of markers such as CDKN2A. Overall, this model for proteinopathy-driven senescence also suggests that photoaged melanocytes may share many features in common with neurons in age-related CNS proteinopathies such as Parkinson’s disease.

These findings also build on previous studies of the role of autophagy in pigmentation by providing evidence that, in addition to contributing to melanosome maturation^54,75^, autophagy may also play a role in the removal of harmful aggregated protein byproducts produced by the tanning pathway. In most cell lineages, autophagy induction is primarily tied to energy sensing through TFEB and TFE3^76,77^. These data support that melanocytes instead regulate autophagy primarily through MITF, ensuring that when the tanning pathway is active, degradative machinery is simultaneously upregulated to maintain proteostasis.

Overall, this study provides evidence that protein misfolding and aggregation are common and physiologically important sequelae of UV exposure. These human and mouse data also point to protein aggregation contributing to common disorders of pigmentation and to melanocyte degeneration with aging, suggesting parallels between the pathophysiology of neurodegenerative disorders and of chronically UV-damaged melanocytes. Future studies may wish to explore targeting of protein aggregates as a therapeutic strategy, as well as explore additional sequelae of proteotoxicity on cell health and organelle viability in the skin.

## Methods

### Cell lines

Newborn foreskin tissues were collected from discarded hospital specimens without any personal identity information. The procedure was approved by the Partners Human Research Committee/IRB (2013P000093). Cells were plated and grown in Medium 254 (Life Technologies #M254500) with Human Melanocyte Growth Supplement (Gibco #S0025). Human melanoma cell line UACC257 (sex unspecified) was obtained from the National Cancer Institute (NCI), Frederick Cancer Division of Cancer Treatment and Diagnosis (DCTD) Tumor Cell Line Repository. SK-MEL-30 (male) human melanoma cell line was from Memorial Sloan Kettering Cancer Center. Both melanoma cell lines have been authenticated by our lab using ATCC’s STR profiling service. UACC257 and SK-MEL-30 cells were cultured in DMEM medium (Life Technologies # 11995073) supplemented with 10% fetal bovine serum and 1% penicillin/streptomycin/L-glutamine in a humidified atmosphere of 95% air and 5% CO2 at 37°C.

### Lentivirus infection

Lentivirus was generated in Lenti-X™ 293T cells (Clontech #NC9834960). The Lenti-X cells were transfected using 250 ng pMD2.G, 1250 ng psPAX2, and 1250 ng lentiviral expression vector in the presence of PEI (MW:25K). For infection with lentivirus, 0.1–1 ml of lentivirus-containing medium was used in the presence of 8 μg/ml polybrene (ThermoFisher #TR1003G). Selection with neomycin (1000 μg/ml) or fluorescence-activated cell sorting for GFP was performed the day after infection. Transduced plasmids include ATG7 CRISPR/Cas9 KO Plasmid (h2) (Santa Cruz, SC-400997-KO-2), pInducer20 (Addgene #44012), and pInducer20^SNCA^ (Addgene #92200).

### Solubility Fractionation Studies

Solubility fractionation of cell lysates was performed as previously published^78^. Briefly cells were grown in monolayer and treated as described. At experimental endpoint, cells were detached with 0.05% trypsin-EDTA (ThermoFisher #25300054) and washed once with PBS before solubilization on ice in 1% triton X-100 in PBS with 1x Halt Protease Inhibitor Cocktail (ThermoFisher #78430). Samples were then transferred to a manual glass dounce tissue grinder for manual disruption. Lysates were then incubated at 4°C with rocking for 30 minutes, at which point samples were freeze-thawed in a –196°C liquid nitrogen bath for 2 minutes followed by 2 minutes of rapid thawing in a 37°C hot water bath for 3 cycles. Samples were then transferred to ultracentrifuge tubes and spun at 100,000g for 30 minutes at 4°C. The supernatant was then transferred to a separate tube and labeled as the triton fraction. Ultracentrifugation pellet was then washed in 100μl of identical triton buffer and ultracentrifuged again. The remaining pellet was solubilized in 2% SDS in 1 M Tris pH 7.4 with halt inhibitor cocktail and labeled as the SDS fraction.

### Immunoblotting

Whole-cell protein lysates were prepared using RIPA lysis buffer (Sigma-Aldrich, #R0278) supplemented with Halt Protease and Phosphatase Inhibitor (ThermoFisher Scientific, #78430). Protein concentrations were quantified using the Pierce BCA protein assay (ThermoFisher #A55864). Immunoblotting was performed by standard techniques using 4-20% Mini-PROTEAN TGX Precast gels (Bio-Rad Laboratories, #4561094) and transferring to 0.45 μm PVDF membranes (Life Technologies, #88518). Membranes were blocked with 5% non-fat milk (Santa Cruz Biotechnology #SC-2325) in TBS containing 0.1% Tween 100 and incubated with one of the following primary antibodies at the indicated dilution: 1:20 dilution of anti-MITF monoclonal antibody C5 (Dana Farber), 1:1000 dilution of anti-alpa-synuclein MJFR1 (abcam #ab138501), 1 µg/mL anti-alpha-synuclein (Syn 211) (Invitrogen #AHB0261), 1:500 anti-PMEL (HMB45) (Invitrogen, #MA134759), 1:3000 anti-vimentin (D21H3) (Cell Signaling #5741), 1:1000 anti-GAPDH (14C10) (Cell Signaling #2118), 1:1000 anti-ATG7 (Cell Signaling #8558), 1:10,000 beta-actin (BA3R), HRP (Invitrogen #MA5-15739-HRP), 1:1000 anti-SQSTM1/p62 (D6M5X) (Cell Signaling #23214T), 1:1000 anti-LC3B (D11) XP (Cell Signaling #3868), 1:1000 anti-gp100 (EP4863(2)) (abcam #ab137078). Incubation with the appropriate secondary antibody followed, either a 1:5,000 dilution of Anti-rabbit IgG, HRP-linked Antibody (Cell Signaling #7074) or a 1:5,000 dilution of anti-mouse IgG, HRP-linked Antibody (Fisher Scientific #50-195-914). To verify equal loading of samples, membranes were re-probed with indicated loading controls. Protein bands were visualized using Immobilon Western Chemiluminescent HRP Substrate (Millipore # WBKLS0) and quantified using ImageJ software (NIH).

### RT-qPCR

Total RNA was isolated from cultured primary melanocytes or melanoma cells using the QIAcube Connect DNA and RNA extraction system (Qiagen #9002864). mRNA expression was determined using intron-spanning primers with SYBR FAST qPCR master mix (Kapa Biosystems, #KK4600). Expression values were calculated using the comparative threshold cycle method (2−ΔΔCt) and normalized to human RPL11 mRNA.

### siRNA transfection

A single treatment of 10 nmol/L of siRNA was delivered to a 60% confluent culture by transfection with lipidoid transfection reagent made by our lab according to a published protocol^79^. After 48-72 hours of transfection, total protein was harvested.

### Cellular melanin quantification

Equal numbers of cells were plated in 6-well plates and treated as described. The cells were then trypsinized, washed in PBS, and split in half for protein and melanin measurement. Half of the cells were used for measurement of protein concentration with the Pierce BCA protein assay (ThermoFisher #A55864) and half were resuspended in 60 μl of 1 N NaOH solution and incubated at 60°C for 2 h or until the melanin was completely dissolved. After cooling down to room temperature, samples were centrifuged at 500 × g for 10 min and the supernatants were loaded onto a 96-well plate. The melanin content was determined by measuring the absorbance at 405 nm on an Envision plate reader, compared with a melanin standard (0 to 50 μg/ml; Sigma Aldrich, #M8631). Melanin content was expressed as micrograms per microgram of protein.

### Mouse strains

Mice harboring the tTA-inducible SNCA^WT^ (B6;D2-Tg(tetO-SNCA)1Cai/J) or SNCA^A53T^ ^(^Tg(tetO-SNCA*A53T)E2Cai/J) transgenes were purchased commercially (Jackson Laboratory strains # 012450 and 012442)^39^ and bred with mice harboring the dopachrome tautomerase-specific response gene (Dct-rtTA, a generous gift of Glenn Merlino, NCI). These strains were also cross-bred with mice harboring the K14-SCF mutation^80^, which produces offspring with interfollicular epidermal melanocytes, recapitulating human epidermal anatomy. ATG7^flox/flox^ mice (C57BL/6J-Atg7em1Lutzy/J) were also commercially purchased (Jackson Laboratory strain # 034429) and bred with mice harboring Cre-recombinase linked to the tyrosinase promoter (Ty-Cre)^62^ for melanocyte-specific ATG7 deletion before breeding onto the base SNCA^A53T^ overexpression genotype. TFEB^flox/flox^ mice (a generous gift of Andrea Ballabio, Fondazione Telethon) were cross-bred with Ty-Cre mice before breeding with the SNCA^A53T^ model. Mouse pups were genotyped using the Transnetyx genotyping service (transnetyx.com). Mouse pups were genotyped using the Transnetyx genotyping service (transnetyx.com). SNCA^A53T^ mice homo– vs. heterozygous for the Dct-rtTA and SNCA overexpression transgenes were verified by backcross to WT B6 mice. Mice used for experiments were all heterozygotes unless noted otherwise. Mice were genotyped through real-time PCR using specific probes designed for each gene (TransnetYX, Cordova, TN).

### Mouse treatments and photography

All animal procedures were conducted in accordance with protocols approved by the Massachusetts General Hospital Institutional Animal Care and Use Committee (IACUC), and animals cared for according to the requirements of the National Research Council’s Guide for the Care and Use of Laboratory Animals (#2008N000025). For benzoyl peroxide treatments (BPO), mice were shaved weekly with Wahl® MiniArco electric trimmers (Patterson Veterinary Supply Inc #07-837-3782). After trimming, 100μl of 10% w/v Luperox A98 Benzoyl Peroxide (ThermoFisher #NC0592884) in acetone or acetone vehicle was applied to the skin of mice three times per week for the indicated number of weeks. For UVA treatments, mice were first trimmed as above and then further depilated with nair hair removal lotion and washed liberally with water once weekly. Mice were then placed in an irradiation chamber and exposed to 5J/cm^2^ UVA radiation (peak em. 368nm).

For photography, mice were anesthetized with inhaled isoflurane anesthesia. Photos were obtained using a Nikon D750 Digital Camera inside a FotodioX LED Studio-In-a-Box (B&H #FOSIAB1616) in standardized fashion.

### Histology and immunostaining studies

For histology, paraffin sections were prepared and stained with hematoxylin and eosin (H&E) or Fontana-Masson stain using the iHisto service (https://www.ihisto.io/). For immunofluorescence, paraffin sections were deparaffinized by xylene and rehydrated gradually with ethanol to distilled water. Sections were submerged in 0.01 M citrate buffer and boiled for 10 min AT 95°C for retrieval of antigen. The sections were washed with PBS-T (0.1% Tween 20) and blocked with 10% normal goat serum (Life Technologies #50062Z) for 1 h at room temperature before application of primary antibody at manufacturers’ recommend concentrations diluted in 50% PBS and 50% goat serum overnight at 4°C. The following day, sections were washed with PBS-T three times and incubated with secondary antibody Goat anti-Rabbit IgG (H+L) Cross-Adsorbed Secondary Antibody, Alexa Fluor™ 568 (Invitrogen #A-11011), Goat Anti-Mouse Alexa Fluor+ 594 (Life Technologies A32742), Goat anti-Rabbit IgG (H+L) Highly Cross-Adsorbed Secondary Antibody, Alexa Fluor™ Plus 488 (Life Technologies #A32731), IgG (H+L) Highly Cross-Adsorbed Goat anti-Rabbit, Alexa Fluor® 647 (Invitrogen #A21245), Goat anti-Mouse IgG (H+L) Highly Cross-Adsorbed Secondary Antibody, Alexa Fluor™ Plus 647 (Life Technologies #A32728), Alexa Fluor® 488 AffiniPure VHH Fragment Alpaca Anti-Mouse IgG (H+L) (Jackson ImmunoResearch #615-544-214). After washing, the tissue sections were cover-slipped with mounting medium (ProLong Diamond Antifade Mountant with or without DAPI, ThermoFisher Scientific, #P36966 or P36965). For thioflavin staining, SYTOX™ Deep Red Nucleic Acid Stain (ThermoFisher #S11380) was used for nuclear visualization. TrueBlack® Plus Lipofuscin Autofluorescence Quencher (ThermoFisher #NC2015600) was used to quench skin tissue autofluorescence according to the reagent instructions.

The following primary antibodies were used at the indicated dilutions: 1:50 anti-PMEL (HMB45) (Invitrogen, #MA134759), 1:25 anti-alpha-synuclein Antibody (LB 509) (Santa Cruz Biotechnology #sc-58480), 1:5000 (human) or 1:1000 (mouse) anti-alpha-synuclein aggregate antibody [MJFR-14-6-4-2] – conformation-specific (abcam #ab209538), 1:100 anti-SOX10 (D5V9L) (Cell Signaling #89356),1:100 anti-Tyrosinase, clone T311 (Sigma #05-647), 1:50 anti-CD163 [OTI2G12] (abcam # ab156769), 1:250 anti-F4/80 (D2S9R) (Cell Signaling #70076).

Primary human melanocytes (10,000-100,000 cells/well) were cultured on Nunc™ Lab-Tek™ II Chamber Slide™ System (ThermoFisher, #154526). After experimental treatments, the cells were fixed with 4% paraformaldehyde (PFA) for 20 min at room temperature, followed by treatment with 0.3% Triton X-100 (Sigma Aldrich #T9284) for 5 min and blocking with 10% goat serum (Life Technologies #50062Z) for 60 min at room temperature. Cells were incubated with the indicated primary antbodies overnight at 4°C. The following day, the slides were washed with PBS-T three times and incubated with secondary antibodies (1:400). Sections were washed with PBS-T three times and mounted in mounting medium (ProLong Diamond Antifade Mountant with or without DAPI, ThermoFisher Scientific, #P36966 or P36965). Thioflavin-stained cells were stained with SYTOX™ Deep Red Nucleic Acid Stain for nuclear visualization.

### Thioflavin stain with immunostaining

After deparaffinization and rehydration, slides were then washed in phosphate-buffered saline (PBS) for 5 minutes, permeabilized in 0.2% triton in PBS for 10 minutes, then washed with 0.01% Tween in PBS twice for 5 minutes each. Antigen retrieval was performed at 95°C for 20 minutes. Following antigen retrieval, the slides were washed with PBS-T then rinsed with PBS briefly. A 0.05% Thioflavin S (Sigma Aldrich #T1892) in 1:1 EtOH:H2O solution was prepared, filtered through a 22-micron filter, and covered with aluminum foil before use. Tissue sections were incubated with Thioflavin S solution at room temperature for 15 minutes covered by aluminum foil. For the remainder of the protocol, slides were covered with aluminum foil to block light exposure. The slides were then washed at room temperature in 50% EtOH twice for 20 minutes each, then in 80% EtOH for 20 minutes, and finally in 0.01% Tween in PBS for 5 minutes. Tissue sections were briefly rinsed with H2O prior to blocking with 10% goat serum for 1 hour at room temperature After blocking, the primary antibodies diluted to the appropriate stock concentration in 50/50 10% goat serum and PBS were added to the tissue sections and left to incubate overnight at 4°C. After wash, tissue sections were then incubated with secondary for 1 hour at room temperature followed by washing with PBS-T. Slides were incubated in SYTOX Deep Red Nucleic Acid Stain (Thermo Fisher #S11380) diluted in PBS 1:2000 for 30 minutes, mounted with Prolonged Diamond antifade reagent (Thermo Fisher #P36965) and a glass cover slip before sealing with nail polish.

### Human biopsies

Human biopsies of pigmentation disorders obtained as part of routine clinical care were obtained from MGH (Hoang), BWH (Murphy), and MGPO dermatopathology (Horn, Lin) as approved by the Mass General Brigham Human Research Affairs Institutional Review Board (IRB# 2020P003834). Human biopsies of pigmentation disorders obtained as part of routine clinical care were obtained from Massachusetts General Hospital (Hoang), Brigham and Women’s Hospital (Murphy), Beth Israel Deaconess Medical Center (Shaw), and Mass General Physicians Organization dermatopathology (Horn, Lin). De-identified patient samples labeled only with a diagnosis were provided to laboratory personnel for staining and quantitation. A minimum of five separate donors for each diagnosis (solar lentigo, melasma, fixed drug eruption, post-inflammatory hyperpigmentation) were stained for dermal synuclein, and variable amounts of synuclein-positive dermal material were identified in all samples. Appropriate positive and negative controls were used for quality control.

### Image quantitation

Images were captured using confocal microscopy (Zeiss Axio Observer Z1 Inverted Phase Contrast Fluorescence microscope (Zeiss) or Nikon A1SiR on a TiE inverted scope (Nikon)). Tissue sections stained with H&E or special stains were digitized using a Nanozoomer S60 (Hamamatsu #C13210-01). Images were processed using NIS-Elements (Nikon) and ZEN Blue (Zeiss) for presentation.

Mouse ear and skin darkness were quantified using ImageJ (version 1.53k). Photographs were converted to 8-bit grayscale and minimum three regions of interest per animal were measured for pixel saturation. Background signal was subtracted before statistical analysis. Photography was standardized as described above.

For quantitation of Sox10-positive melanocytes or F4/80-postive macrophages stained by immunohistochemistry, five high-powered fields per sample were chosen at random. De-identified images were given to authors uninvolved in image collection (Ji, Joseph) and positive cells per field were quantified. For melanophage quantitation, cells which were showed evidence of both F4/80 staining and melanin were quantified.

Punctae in UV-treated melanocytes were measured for a minimum size as described above and quantified in similar fashion. Five high-powered fields for each sample were again utilized.

### Mouse fur melanin quantitation

As previously described^81^, dorsal hairs of mice at 8 weeks of age were shaved and 1 mg was dissolved overnight in 1 mL of 9:1 Soluene-350 (Perkin Elmer #6003038) and water. Triplicate 150 μL aliquots for each mouse hair sample were then analyzed for absorbance values at 405 nm.

### Immunoprecipitation

Immunoprecipitation for mass spectrometry was performed using SK-MEL-30 melanoma cells transduced with the pInducer20^SNCA^ plasmid for doxycycline-inducible SNCA overexpression in order to promote aggregation and provide sufficient SNCA yield for analysis or empty backbone. Briefly, cells were treated as described before being lysed in buffer containing 1% triton X-100 and PBS with halt protease and phosphatase inhibitor cocktail. Lysates were pre-cleared for 1 hour using protein A-linked dynabeads (Life Technologies # 10001D). Supernatant was then isolated and incubated overnight at 1:600 dilution of MJFR1 anti-alpha-synuclein antibody (abcam # ab138501) at 4°C with rotation. The next day, 100 microliters of protein A dynabeads were added to each sample and allowed to incubate with rotation at 4°C for another 4 hours. Samples were then washed five times with 50mM Tris wash buffer before addition of 100mM Tris storage buffer and shipment on dry ice for mass spectrometry analysis.

### Mass spectrometry-based affinity proteomics

#### On-bead trypsin digestion of biotinylated proteins for proteomics

Samples with biotinylated proteins bound to streptavidin magnetic beads were initially washed twice with 200 uL of 50 mM Tris-HCl (pH=7.5) buffer. Proteins were then digested by adding 80uL of the digestion buffer (2M Urea, 50nM Tris-HCl, 1mM DTT, and 0.4ug trypsin) and incubating for 1 hour at room temperature (RT) while shaking at 1000 rpm. The supernatant was collected into a fresh tube, and another 80 uL of the digestion buffer was added for incubation for another 30 minutes. The second supernatant was collected, and beads were washed twice with 60 uL of 2M Urea/50mM Tris-HCl buffer. These washes were combined with the digestion supernatant. The eluate of on-bead digestion was reduced with 4 mM DTT for 30 minutes before being alkylated with 10 mM Iodoacetamide for 45 minutes in the dark, both at RT while shaking at 1000 rpm. Finally, each sample was digested overnight with 0.5ug of trypsin on a Thermo shaker at RT and 1000rpm shaking. Next morning, digested peptides were acidified with neat formic acid (FA) to 1% FA final (pH<3) for desalting on in-house packed C18 (3M) StageTips. Briefly, C18 StageTips were conditioned sequentially with 100 uL of 100% methanol (MeOH), 100 uL of 50% acetonitrile (MeCN)/0.1% FA, and 2x 100uL of 0.1% FA. Peptide samples were loaded onto the C18 StageTips, washed twice with 100 uL of 0.1% FA, eluted from the C18 resin with 50 uL of 50% MeCN/ 0.1% FA, and vacuum-centrifuged until dry.

### TMT labeling and StageTip peptide fractionation

Peptide samples were labeled with TMT6 reagents (Thermo Fisher Scientific). Dry peptide peptides were resuspended in 80 μL of 50 mM HEPES and incubated with 20 uL of the 25 ug/uL TMT6 reagents in MeCN for 1 hour at RT while shaking at 1000 rpm. Next, 4 μL of 5% hydroxylamine was added to quench TMT reaction in each sample, incubated for 15 minutes at RT while shaking. All labeled peptide samples were then combined, vacuum-centrifuged to dry, and desalted on a C18 StageTips using the previously described protocol. The dry combined peptide sample was resuspended in 200 uL of 0.1% FA for basic reverse phase (bRP) fractionation on an in-house packed SDB-RPS (3M) StageTip. Briefly, the SDB-RPS StageTip was conditioned sequentially with 100 μL of 100% MeOH, 100 μL 50% MeCN/0.1% FA, and 2x 100 μL 0.1% FA. The sample was loaded on the conditioned StageTip and then eluted in eight fractions using 20 mM ammonium formate buffers with increasing (vol/vol) concentrations of MeCN (5%, 7.5%, 10%, 12.5%, 15%, 20%, 25%, and 45%). All fractions were vacuum-centrifuged until completely dry.

### Liquid chromatography and mass spectrometry

Peptide fractions were resuspended in 9 μL of 3% MeCN/0.1% FA for analysis on an online liquid chromatography tandem mass spectrometry (LC-MS/MS) system, including a Vanquish Neo UPHLC (Thermo Fisher Scientific) coupled to an Orbitrap Exploris 480 (Thermo Fisher Scientific). Samples were injected onto a microcapillary column (Picofrit with 10 µm tip opening/75 µm diameter, New Objective, PF360-75-10-N-5) packed in-housed to 30 cm with C18 silica material (1.5 µm ReproSil-Pur C18-AQ medium, Dr. Maisch GmbH, r119.aq) and heated to 50 °C using column heater sleeves (PhoenixST). Solvent A (0.1% FA) and solvent B (90% MeCN/0.1% FA) were used for analysis on the LC-MS/MS system. For each peptide fraction, 4 uL was injected into the MS and analyzed on a 110 min method with the following gradient profile (min:%B): 0:1.8; 1:5.4; 85:27; 94:54; 95:81; 109:81 at a flow rate of 200 nL/min. MS operated in data-dependent acquisition mode with MS1 spectra measured at 60,000 resolution, 300% normalized AGC target, and m/z range from 350 to 1800. MS2 spectra of the top 20 most abundant ions per cycle were acquired at 15,000 resolution, 30% AGC target, 0.7 m/z isolation window, and 34 normalized collision energy. The dynamic exclusion time was set to 20 s, and the peptide match and isotope exclusion functions were enabled.

### Mass spectrometry data processing

Mass spectrometry data were processed using Spectrum Mill (proteomics.broadinstitute.org). Spectra within a precursor mass range of 600-6000 Da and a minimum MS1 signal-to-noise ratio of 25 were retained. MS1 spectra within a retention time range of +/− 45 s and a precursor m/z tolerance of +/− 1.4 m/z were merged. Spectra were searched against a human Uniprot database with 602 common laboratory contaminants added. Digestion parameters were set to “trypsin allow P”, and up to 4 missed cleavages were allowed. Fixed modifications included carbamidomethylation on cysteine, TMTPro on the N-terminus and internal lysine. Variable modifications were set as acetylation of the protein N-terminus, oxidation of methionine, N-term Q-pyroglutamate formation, and N-term deamidation. A minimum matched peak intensity of 30%, precursor and product mass tolerance of +/− 20 ppm, and a maximum ambiguous precursor charge of 3 were used as match tolerances. A maximum false discovery rate (FDR) threshold of 1.2% for precursor charges ranging from +2 to +6 was used to validate peptide spectrum matches (PSMs). A target protein score of 0 was applied to further filter PSMs. TMTpro reporter ion intensities were corrected for isotopic impurities using the afRICA correction method, which utilizes determinant calculations according to Cramer’s Rule.

All non-human proteins were removed before statistical analysis on the Proteomics Toolset for Integrative Data Analysis (Protigy, v1.0.7, Broad Institute, https://github.com/broadinstitute/protigy). Each protein was associated with a log2-transformed ratio of every TMT condition to the median intensity of all channels. Protein data were normalized to the median within two separate groups: SNCA-overexpress and SNCA-knockout. After normalization, an empirical Bayes-moderated t-test was used to compare treatment groups using the limma R package. P-values associated with every modified peptide or protein were adjusted using the Benjamini–Hochberg FDR approach.

### Tissue explants

Skin samples considered surgical waste were obtained de-identified from healthy donors (IRB# 2013P000093) undergoing reconstructive surgery, according to institutional regulations. Full thickness human abdominal skin explants were cultured in petri dishes in Williams E media lacking phenol red (Life Technologies #A1217601) and containing 10 vg/ml of insulin (Santa Cruz Biotechnology #SC-29062), 10 ng/ml of hydrocortisone (Sigma Aldrich # H0135) and Penicillin-Streptomycin (Life Technologies #15140163) changed daily. For UV irradiation experiments, a UV lamp (UV Products) was used at 5J/cm2 UVA (peak em. 368nm).

### Transmission electron microscopy

Mouse skin samples were immersion fixed with half strength Karnovsky’s fixative (2% formaldehyde + 2.5% glutaraldehyde in 0.1 M sodium phosphate buffer, pH 7.4; Electron Microscopy Sciences, Hatfield, Pennsylvania) at room temperature then stored under refrigeration. After fixation, samples were rinsed with 0.1M sodium cacodylate buffer, post-fixed with 2% osmium tetroxide in 0.1M sodium cacodylate buffer for 1.5 hours, en bloc stained with 2% aqueous uranyl acetate for 30 minutes, then dehydrated with graded ethyl alcohol solutions, transitioned with propylene oxide and resin infiltrated in EMBED812 epoxy resin (Electron Microscopy Sciences, Hatfield, Pennsylvania) utilizing an automated EMS Lynx 2 EM tissue processor (Electron Microscopy Sciences, Hatfield, Pennsylvania.) The processed samples were oriented into EMBED812 epoxy resin inside flat silicone molds and polymerized in a 65oC oven for 48 hours. Semi-thin sections were cut at 1 μm thickness and stained with 1% toluidine blue in 1% sodium tetraborate aqueous solution and Permount coverslipped on glass slides then scanned using a Hamamatsu Nanozoomer (Hamamatsu Photonics, Bridgewater, NJ) for assessment and ultrathin region-of-interest localization. Ultrathin sections (∼80 nm) were cut from each sample block using a Leica EM UC7 ultramicrotome (Leica Microsystems, Buffalo Grove, IL, USA) and a Diatome ULTRA diamond knife (Diatome, Hatfield, Pennsylvania), then collected using a loop tool onto either 2×1 mm, single slot formvar-carbon coated or 200 mesh uncoated copper grids (Electron Microscopy Sciences, Hatfield, Pennsylvania) and air-dried. The ultathin sections on grids were stained with aqueous 2% uranyl acetate and Sato’s lead citrate stains using a modified Hiraoka grid staining system (Electron Microscopy Sciences, Hatfield, Pennsylvania). Grids were imaged using a FEI Tecnai G2 Spirit transmission electron microscope (FEI, Hillsboro, Oregon) at 80 kV interfaced with an AMT XR41 digital CCD camera (Advanced Microscopy Techniques, Woburn, Massachusetts) for digital TIFF file image acquisition. TEM imaging of the assessed and digital images captured at 2kx2k pixel, 16-bit resolution.

### Flow cytometry

Proteostat staining and flow cytometry was performed using the Proteostat aggresome detection kit (Enzo Life Sciences, Farmingdale, NY, USA) per manufacturer instructions. Cells were treated as described, trypsinized, washed with PBS, and fixed in 4% PFA for 30 minutes at RT. Cells were then washed in PBS and permeabilized using 0.05% triton X-100 for 30 minutes on ice. Cell suspensions were stained using a 1:10,000 dilution of proteostat dye in manufacturer-provided staining buffer and analyzed using a Cytek Aurora Spectral flow cytometer or a LSR Fortessa X-20 flow cytometer. Whole cells were gated based on forward and side scatter, and doublet discrimination was performed. Differences in median fluorescent intensity of proteostat signal were analyzed to determined per-cell misfolded protein load.

### Split luciferase reporter assay

To measure SNCA aggregation, UACC257 melanoma cell lines were infected with split SNCA Luci-ferase reporter system (a generous gift of the Hyman Lab), which expresses alpha-synuclein linked to one of two fragments of Gaussia luciferase (GLuc)^82^. Cells were also infected with either ATG7 CRISPR/Cas9 KO Plasmid (h2) (Santa Cruz, SC-400997-KO-2) or empty vector. Resulting lines were treated with 100nm rotenone as described above for 24 hours and either harvested or washed with PBS and given fresh DMEM media for an additional 24 hours. Luciferase signal was read after addition of colenterazine (ThermoFisher # # 501365238) to a final concentation of 5μM and read immediately using an Envision plate reader at 480nm.

### Processing of human CUT&RUN data

We aligned the raw sequencing reads to HG38 using the Bowtie2 package (version 2.4.5, parameters –-end-to-end –-very-sensitive –-no-mixed –-no-discordant –-phred33 –I 10 –X 700 – dovetail)^83^. We called the binding location of transcription factors using SEACR (1.3) with default parameters^84^.

### Processing of primary melanocyte bulk RNA-seq data

We aligned the raw sequencing reads to HG38 using the HISAT2 package (2.2.1) with default parameters^85^. We quantified gene counts for the UCSC gene annotation using StringTie^86^ and performed differential gene expression analysis between MITF KD, TFE3 KD, and TFEB KD and their respective control samples using the DESeq2 package (1.20.0) with default parameters^87^. We adjusted p-values for multiple testing using the Benjamin-Hochberg method.

### Real-Time Quaking-Induced Converstion Assay (RT-QuIC)

RT-QuIC was performed as previously described^27,28,88^. Briefly, reactions were performed in Nunc black 96-well flat bottom plates (Thermo Fisher). Prior to adding reaction buffer, each well was loaded with three 0.8 mm low-binding silica beads (OPS Diagnostics, Lebanon, NJ, USA). Recombinant SNCA monomer (rPeptide #S-1001) was reconstituted in HPLC-grade water to 1 mg/ml and filtered through Amicon 100 kDa filters (Millipore) by centrifugation for 15 min at 4 °C at 15,000 rcf. Tissue homogenate or cell lysate was diluted as specified, and 2 μl total volume was added to individual wells containing 98 μl of RT-QuIC reaction mixture composed of 40 mM NaPO4 (pH 8.0), 170 mM NaCl, 20 µM ThT (Thermo Fisher), and 0.1 mg/ml SNCA monomer. The plates were sealed with Nunc clear sealing film (Thermo Fisher) and incubated at 42 °C in a BMG FLUOstar Omega plate reader (BMG Labtech) with cycles of 1 min shaking (400 rpm, double orbital) and 1 min rest throughout the assay. ThT fluorescence (450 nm excitation and 480 nm emission; bottom read) was recorded every 30 min for 72 hours. Four replicate reactions were made for each sample. The plate reader was set up with optic gain setting (∼ 1900) that would give a negative control baseline of ThT fluorescence around 15,000-25,000 relative fluorescence units (rfu). Where applicable, a “positive” result from an individual well was defined as ThT fluorescence greater than 50,000 out of a maximum value of 270,000 rfu. Samples were analyzed both for average fluorescence across four technical replicates for a given samples at a given dilution and for total number of “positive’ wells by 72 hour endpoint.

### Preparation and use of SNCA pre-formed fibrils (PFFs)

Recombinant monomeric SNCA was purchased commercially (rPeptide #S-1001) and resuspended to a final concentration of 5mg/ml in sterile PBS. Suspensions were then incubated for seven days at 37°C at 1000rpm in a MultiTherm Shaker (Benchmark Scientific Inc. #H5000-H) until suspensions became turbid. PFF formation was verified in 96 well format using thioflavin T dye fluorescence emission at 485nm. Before transfection, PFFs were sonicated using a Fisherbrand Model 120 Sonic Dismembrator for 60 seconds per sample in pulses of 1 second on, 1 second off at 30% amplitude. α-Synuclein pre-formed fibrils (PFFs) were adsorbed onto 400-mesh carbon-coated copper grids (Electron Microscopy Sciences, cat. # FCF400-CU) for 1 min, then stained by floating the grids on 10 µL of 2% uranyl acetate for 2 minutes to ensure appropriate fibrils size. After a quick wash with distilled water, the grids were air-dried prior to imaging on a JEOL JEM-1011 transmission electron microscope with AMTv601 software (Advanced Microscopy Techniques, Woburn, MA, United States). Following sonication, the mean fibril length was approximately 50 nm.

### Mouse skin preparation for scRNA-seq

Mice were euthanized via CO_2_ inhalation until breathing and movement ceased. Flank skin was then removed using sterile scissors and incubated in dispase at 37°C for 90 minutes. Epidermis was peeled from dermis and transferred to 0.25% trypsin-EDTA solution where it was minced with #15 scalpels and incubated with rocking for 20 minutes at 37°C. Cell suspensions were then serially filtered through 100, 70, and 40 μm cell stainers (Thomas Scientific, Logan Township, NJ, USA), washed in PBS, and depleted of dead cells using the EasySep Dead Cell Removal (Annexin V) Kit (StemCell Technologies Inc. Vancouver, Canada). Live cells were then counted and submitted for single cell RNA sequencing.

### 10X Single Cell V3.1 3’ Feature Barcoding Method

Cell suspensions are handed off to the Broad Clinical Labs Single Cell processing team. Appropriate volume adjustments are made to ensure the target cell recovery is met prior to the addition of reverse transcription mastermix. The cells in the mastermix are then added to the 10x Chip G along with 10x gel beads and partitioning oil. The chip is run on the 10x Chromium Controller to create an emulsion of single cells, RT mastermix, and a barcoded gel bead. After emulsions are created, samples are immediately put on reverse transcription thermalcycler protocol set at 53OC for 45 minutes, 85OC for 5 minutes before holding at 4OC. The emulsion is then broken, and debris is removed via a Silane bead cleanup. The purified cDNA is then amplified with the following PCR conditions: Initial denaturation at 98°C for 3 minutes, followed by 12 cycles of denaturation at 98°C for 15 seconds, annealing at 63°C for 20 seconds, and extension at 72°C for 1 minute. A 0.6X SPRI cleanup is performed to generate gene expression (GEX) and feature barcoding (FBC) cDNA before being quantified on a Qubit.

GEX cDNA library construction is run on an entirely automated workflow. The GEX cDNA is fragmented and repaired with the addition of an A-tail at 32OC for 5 minutes then 65OC for 30 minutes. A 0.8X double sided SPRI cleanup then removes undesired fragments. Adapters are ligated onto the DNA at 20OC for 15 minutes followed by a 0.8X SPRI. 10x dual indexed barcodes are added before a PCR reaction with the following conditions: An initial denaturation at 98°C for 45 seconds, followed by 12 cycles of denaturation at 98°C for 20 seconds, annealing at 54°C for 30 seconds, and extension at 72°C for 20 seconds. These libraries undergo a final 0.8X double-sided SPRI cleanup for size selection. FBC cDNA immediately began with a PCR reaction with 10x dual-indexed barcodes, with the above conditions, but repeating 10 cycles instead. Following PCR Cleanup, quality is confirmed using Quant-iT PicoGreen and an Agilent TapeStation to confirm library size. Samples are normalized to 2nM and pooled together to accommodate the desired reads per cell.

### Illumina Sequencing

Pooled libraries were denatured using 0.1 N NaOH prior to sequencing. Flowcell cluster amplification and sequencing were performed according to the manufacturer’s protocols using either the NovaSeq. Libraries were sequenced with a read structure of 28×10×10×90 using the Broad Clinical Labs Walkup Sequencing product. Data was delivered in the form of FASTQ files by The Broad Institute’s Genomics Platform.

### Processing of mouse scRNA-seq data

The 10x Genomics scRNA-seq data were aligned to mm10 using Cellranger (7.1.0)^83^ and analyzed using the Seurat package (5.3.0)^84^. To eliminate doublets and non-viable cells, the dataset underwent several initial preprocessing steps. Genes with fewer than three UMI, as well as cells with either fewer than 200 or more than 6,000 detected genes, were excluded from the analysis. The subsequent analysis thus included only those genes which satisfied the constraints nCount_RNA ≥ 3 and 200 ≤ mFeature_RNA ≤ 6000, respectively. Furthermore, cells in which 15% or more of the cell’s total UMI counts mapped to mitochondrial DNA were excluded from downstream analysis. The preprocessed count data were normalized for sequencing depth using the NormalizeData function with default parameters (normalization.method = “LogNormalize”, scale.factor = 10000). Based on the normalized dataset, the 2,000 most variable features were identified and subsequently used for principal component analysis (PCA). The top 30 principal components were then utilized for clustering and visualization using Uniform Manifold Approximation and Projection (UMAP). Cluster identification was carried out using established biomarkers from singleCellBase^2^.

The cluster exhibiting the highest expression levels of the melanocyte markers *Mitf*, *Tyr*, *Dct*, *Mlana*, *Tyrp1*, *Kit*, and *Sox10* was designated as the melanocyte cluster. An adjacent cluster, located near the melanocyte population on the UMAP plot and characterized by high expression of both melanocyte markers and keratinocyte-associated genes (*Cdh1*, *Krt5*, *Krt10*, *Krt14*, and *Trp63*), was identified as a population of melanocyte-keratinocyte doublets. This doublet cluster was excluded from all subsequent downstream analyses.

Using Seurat’s AverageExpression method with default parameters, we estimated that within the melanocyte cluster overexpressed SNCA levels were approximately 3 times larger than basal Snca in both the hyper and hypo groups.

### Differential expression analysis

Differential gene expression analysis was conducted exclusively within the melanocyte cluster using the Wilcoxon Rank Sum test on log-normalized expression data. Comparisons were made between the a53t-hyperpigmented and the k14-control groups (hyper vs. control), the a53t-hypopigmented and the k14-control groups (hypo vs. control), or the a53t-hypopigmented and the a53t-hyperpigmented groups (hypo vs. hyper). Genes expressed in fewer than 1% of cells or exhibiting an average log_10_ fold change of less than 0.1 were excluded from the analysis. Statistical significance was determined using a Bonferroni adjusted *p*-value threshold of *p* < 10^−3^. To investigate the impact of *SNCA* overexpression in the hyperpigmented and hypopigmented groups, six gene sets were curated, corresponding to the following biological categories: pigmentation, neurodegeneration, lysosome, autophagy, cellular senescence, and the unfolded protein response (UPR). These gene sets were used for the generation of heat maps with unbiased hierarchical clustering. Each set was constructed by taking the union of relevant genes from Gene Ontology (GO) categories and Kyoto Encyclopedia of Genes and Genomes (KEGG) pathways. In the case of the neurodegeneration category, all Parkinson’s-implicated genes from the Gene4PD database^3^ were used. The specific GO accession numbers and KEGG pathway identifiers used for each category are provided in the Supplementary Table 1-3.

For the construction of heat maps, genes for which Seurat’s pseudobulk expression values were below 10^#%^ across all experimental groups were considered not expressed within the melanocyte cluster and excluded from further analysis. To generate coarse-grained expression profiles, cells from each group were randomly partitioned into ten blocks. The average gene expression within each block was calculated using log-normalized counts. This process yielded 30 aggregated expression profiles (10 blocks per group: control, hyperpigmented, and hypopigmented), which were subjected to unsupervised hierarchical clustering using an average linkage method with Euclidean distance. This was carried out in an unbiased manner by including all genes belonging to a curated category. In every case, clustering exactly partitioned coarse-grained gene profiles according to their experimental groups with no intermixing, highlighting the reproducibility of the data. For display purpose, the top (up to 15) differentially expressed genes per category first were selected based on the highest absolute average log_2_ fold change values, and the selected genes were organized into upregulated and downregulated groups; finally, only those genes demonstrating statistical significance with a Bonferroni adjusted *p*-value < 10^−3^ in either the hyper vs. control or hypo vs. control comparisons were included in the heat maps. All heat map blocks were rank-ordered and scaled to the interval [0, 1], thereby enabling a uniform distribution of values across the color scale to enhance visual interpretability of expression differences.

### Gene Ontology Analysis

Lists of differentially expressed genes (DEGs) for hyper vs. control and hypo vs. control were analyzed for functional enrichment using the Database for Annotation, Visualization, and Integrated Discovery (DAVID)^4^. DAVID’s functional annotation clustering tool was utilized to identify biological pathways and processes associated with each gene set. Results were visualized using scatter plots in which the negative log_10_ Benjamini-Hochberg adjusted *p*-value was plotted against the enrichment score of each functional annotation cluster.

### Human IGH Samples

Patient volunteers were recruited from general dermatology clinics at Massachusetts General Hospital as approved by the Mass General Brigham Human Research Affairs Institutional Review Board (IRB #2024P000545) Presence of lesions clinically consistent with IGH were verified by a board-certified dermatologist (NT) at time of collection. Three IGH lesions were identified and photographed, as were perilesional skin sites approximately 5 cm away from each lesion. Lesions and perilesional skin were prepped with topical EtOH and anesthetized with 1% lidocaine without epinephrine. Samples were obtained using the shave technique with a dermablade (Accutec Blades, Inc. Verona, VA, USA) and kept in sterile PBS on ice for immediate processing for single cell RNA sequencing identical to mouse flank skin as described above. Each sample was divided in half, with one half used for single cell dissociation and the other fixed in formalin and embedded in paraffin for diagnostic verification by a board-certified dermatopathologist (T.H.)

### Processing of IGH scRNA-seq Data

The 10x Genomics scRNA-seq analysis was performed with the R Seurat package (v5.2.0)^1^. Data from eight samples corresponding to lesional and perilesional shaves for four human patients were analyzed, yielding four biological replicates. Patients 1 and 2 were male; 3 and 4 were female. Identical standard preprocessing steps were applied uniformly to all eight samples: removing genes with fewer than three UMI, removing cells with fewer than 200 or more than 6000 detected genes, and further removing cells in which 10% or more of the cell’s total UMI counts mapped to mitochondrial DNA. Respectively, these filtering steps correspond to the inclusion of data satisfying the constraints nCount_RNA ≥ 3, 200 ≤ mFeature_RNA ≤ 6000, and percent_MT < 10. The filtered data were then normalized using Seurat’s NormalizeData function with default parameters (normalization.method = “LogNormalize” and scale.factor = 10000). Variable features were identified and scaled using Seurat’s FindVariableFeatures and ScaleFeatures, respectively, using the default parameters.

After preprocessing, all samples were integrated using the Canonical Correlation Analysis (CCA) method (Seurat’s IntegrateLayers function with “method = CCAIntegration”). Principal component analysis (PCA) was performed, and the resulting 20 largest PCA dimensions were used for clustering (FindClusters with default arguments and a clustering resolution of 0.2). The Uniform Manifold Approximation and Projection (UMAP) was used to visualize the resulting clusters (Extended data Fig. 9b). A dot plot with cell-type biomarkers was constructed and used for cluster identification (Extended data Fig. 9c). The choice of included biomarkers for the dot plot were taken from manually curated studies compiled in the singleCellBase database^2^. In particular, the biomarkers *MITF*, *TYR*, *DCT*, *MLANA*, *TYRP1*, *PAX3*, *KIT*, and *SNCA* were used to identify the melanocyte cluster. To remove potential doublets and keratinocyte contamination, the melanocyte cluster was subjected to further subclustering using Seurat’s FindSubCluster function (with argument “resolution = 0.1”). This split the melanocyte cluster into two subclusters which were identified as pure melanocytes (cluster 2_0) and multiplets (cluster 2_1). A scatter plot of mean melanocyte versus mean keratinocyte biomarker expression across all four patients (lesional and perilesional combined) was used to confirm the subclustering. Only the pure melanocyte cluster was used for downstream analysis.

### Differential Expression Analysis of the IGH Patient Data

Differential expression analysis was conducted between paired lesional (case) and perilesional (control) samples for each of the four patients using Seurat’s FindConservedMarkers wrapper for the FindMarkers function with default parameters. The resulting patient-specific *p*-values for each gene were then combined and adjusted using Stouffer’s Z-score method and Benjamini-Hochberg adjustment, respectively. The top 300 genes with largest (patient) mean average log_&_ fold-change (lesional/perilesional) and *p*_’()_. < 10^−3^ were taken as the UP gene set. Similarly, the DOWN gene set included the 300 genes with smallest (patient) mean average log_&_ fold-change and *p*_’()_. < 10^−3^.

### Gene Ontology Analysis of the IGH Patient Data

Functional annotation clustering of the differentially expressed UP and DOWN gene sets was performed using DAVID^2^. Expression fold-changes of DE genes from selected UP and DOWN Gene Ontology (GO) categories of biological interest (Supplementary Table 1-3) were displayed as a heatmap (Extended data Fig. 9e). In Fig. 9f, the same combined adjusted p-value threshold was employed, but all genes satisfying the condition of absolute average log_2_ FC > 0.2 were used for GO analysis.

### Analysis of Overlapping DE Genes in SNCA-overexpressed Mouse and IGH Patient Data

The R package biomaRt was used to determine human gene orthologs of genes present in the SNCA-overexpressed mouse analysis. Each mouse and human “mart” was built using biomaRt’s useMart function with arguments ENSEMBL_MART_ENSEMBL; dataset = “mmusculus_gene_ensembl” or “hsapiens_gene_enseml”, respectively; and host = https://dec2021.archive.ensembl.org. The two marts were then linked for homology mapping using getLDS.

## Statistical analysis

Statistical analyses were performed using GraphPad Prism 10. In general. for comparisons of two groups, significance was determined by two-tailed, unpaired Student’s t tests. The specific statistical tests used for experiments are described in the figure legends. P values less than 0.05 were considered statistically significant. Levels of significance are indicated by *p<0.05, **p<0.01, ***p<0.001, ****p<0.0001; ns, not significant in cases where exact p values are not labeled.

## Acknowledgments

The authors would like to thank Todd Ridky, MD, PhD (U. Penn), John Seykora, MD, PhD (U. Penn), and Bruce Spiegelman, PhD (Dana Farber Cancer Institute) for their input regarding conceptualization and study design, as well as Dexter Dean (Georgia Tech) and Kenneth Rose (MGH) for their technical assistance with the RT-QuIC assay. We thank Jennifer C. Lee (National Heart, Lung, and Blood Institute, NIH) for the gift of purified PMEL RPT.

P.B.H, M.H. and J.S.S. were supported in part by NIH R01CA163336. D.E.F. acknowledges support from NIH grants P01 CA163222, R01 AR072304, and R01 AR043369. Department of Defense Melanoma Academy (W81XWH2220052), the Melanoma Research Alliance, Dr. Miriam and Sheldon G. Adelson Medical Research Foundation, the Lancer Professorship in Dermatology (Harvard Medical School) the Water Cove Charitable Foundation. The Sheldon Adelson Medical Foundation also supported the work of N.H., S.A.C, and N.D.U. B.T.H. acknowledges support from The Massachusetts Alzheimer Disease Research Center P30AG062421. N.T. was supported by a Career Development Award from the Dermatology Foundation, as well as the Harvard Dermatology Training Grant (5T32AR007098-51) and by funding from the American Skin Association Mulvaney Award. Transmission electron microscopy studies performed by P.S. were supported in part through NIH National Eye Institute Core Grant P30EY003790. Synuclein knockout mouse breeding was supported by The Nina Compagnon Hirshfield Parkinson’s Disease Research Fund (S.S.C.). A.B. was funded by the European Research Council Advanced Grant (ERC-AdG) INherited CANcer TARgeting (INCANTAR), NIH RO1 grant (R01CA260205-01A1), Italian Telethon Foundation, and Italian Association for Cancer Research (IG-IG 29230). Figure 6a and Extended Data Figure 10 were made using BioRender.com.

## Contributions

N.T and D.E.F. conceived of the study and designed experiments with additional input from B.T.H, A.K.G., J.C.L., and R.B. N.T. performed most bench experiments and analysis. P.B.H. and M.H. performed RNA sequencing analysis supervised by J.S.S with assistance from N.H. Bench experiments and mouse colony maintenance were also performed by T.A.C., R.A,J., H.J., A.E.G, J.R.B., M.Y.D., A.A., and A.M. P.S. performed TEM sample preparation and assisted with visualization. S.M.O, K.K., and Y.Z. assisted with human sample procurement and processing, as well as mouse skin dissociation for RNA sequencing. A.F. assisted with RT-QuIC and SNCA PFF studies as well as PFF quality control. Procurement and IRB protocol management was overseen by S.G. Human clinical sample procurement was overseen by T.D.H., G.F.M., K.C.S., W.L., and M.P.H. Human primary cell harvesting was done with assistance from X.W. Knockout mouse studies were performed with additional technical input from S.S.C. and A.B. Mass spectrometry experiments and downstream analysis were performed by K.N. overseen by N.D.U. and S.A.C. Manuscript text and figures were assembled by N.T. overseen by D.E.F.

Correspondence and requests for materials should be addressed to Corresponding authors David Fisher and Nicholas Theodosakis

---

**Extended Data Fig. 1:**
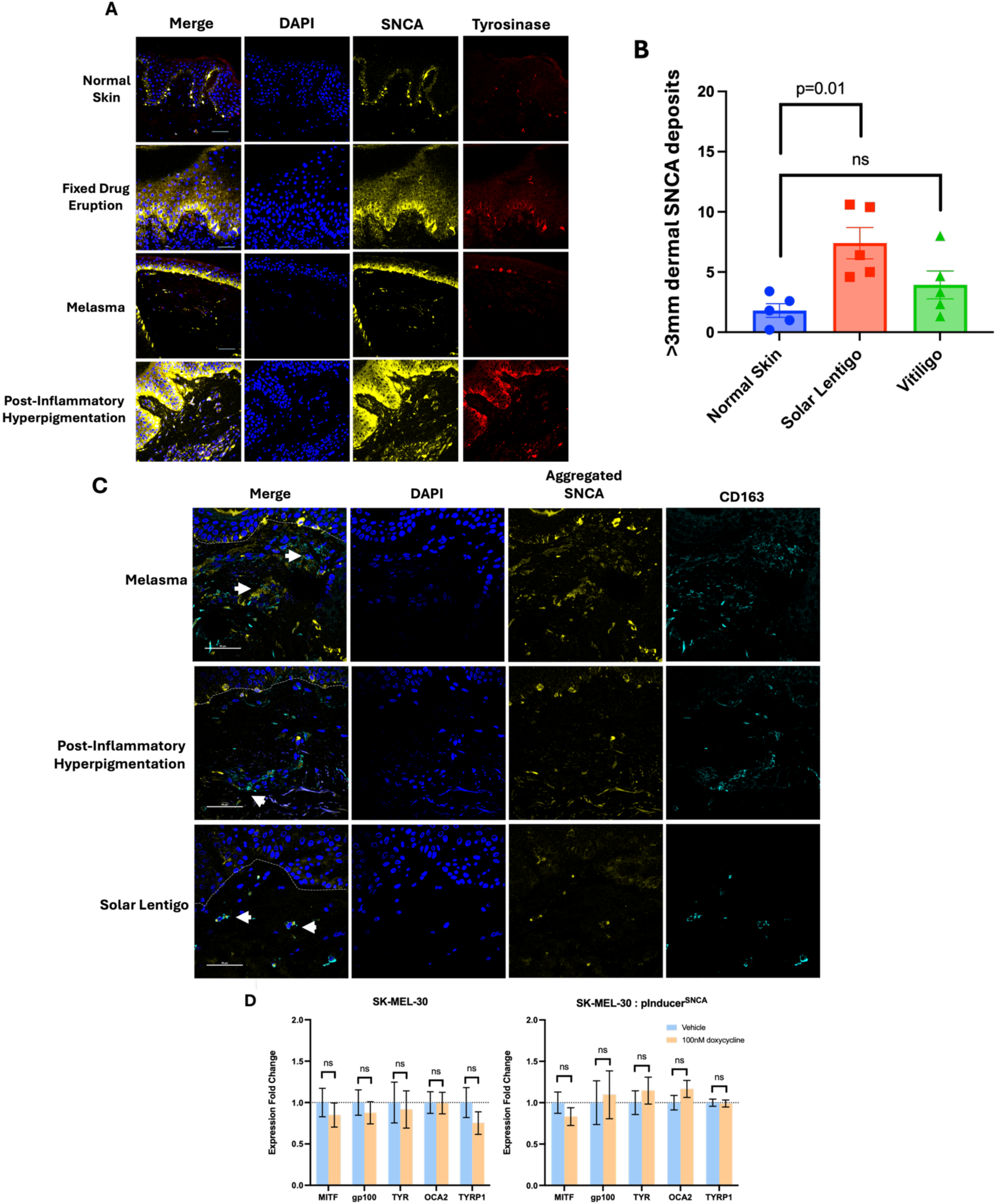
Alpha-synuclein is enriched in hyperpigmented human skin and co-localizes with other melanosomal proteins. (A) Immunofluorescent staining of biopsies of a variety of hyperpigmenting disorders demonstrating colocalization and enrichment of non-aggregated as well as aggregated SNCA both within and around epidermal melanocytes and in the dermis. Normally-pigmented human abdominal skin is shown for comparison. (B) Quantification of dermal SNCA shows enrichment in a cohort of solar lentigines vs normal skin or vitiligo (n=5 donors per condition, student’s *t*-test). (C) Dermal macrophages in a range of hyperpigmenting disorders are frequently found in close proximity to extracellular aggregated SNCA in the dermis (arrowheads). Scale bar = 50μm in all images, DEJ are labeled with a white dashed line. (D) 24 hour overexpression of SNCA in SK-Mel-30 melanoma cells stably transduced with the pInducer20^SNCA^ plasmid suggests synuclein levels do not directly induce changes in MITF target gene expression (student’s *t*-test, n=5).

**Extended Data Fig. 2:**
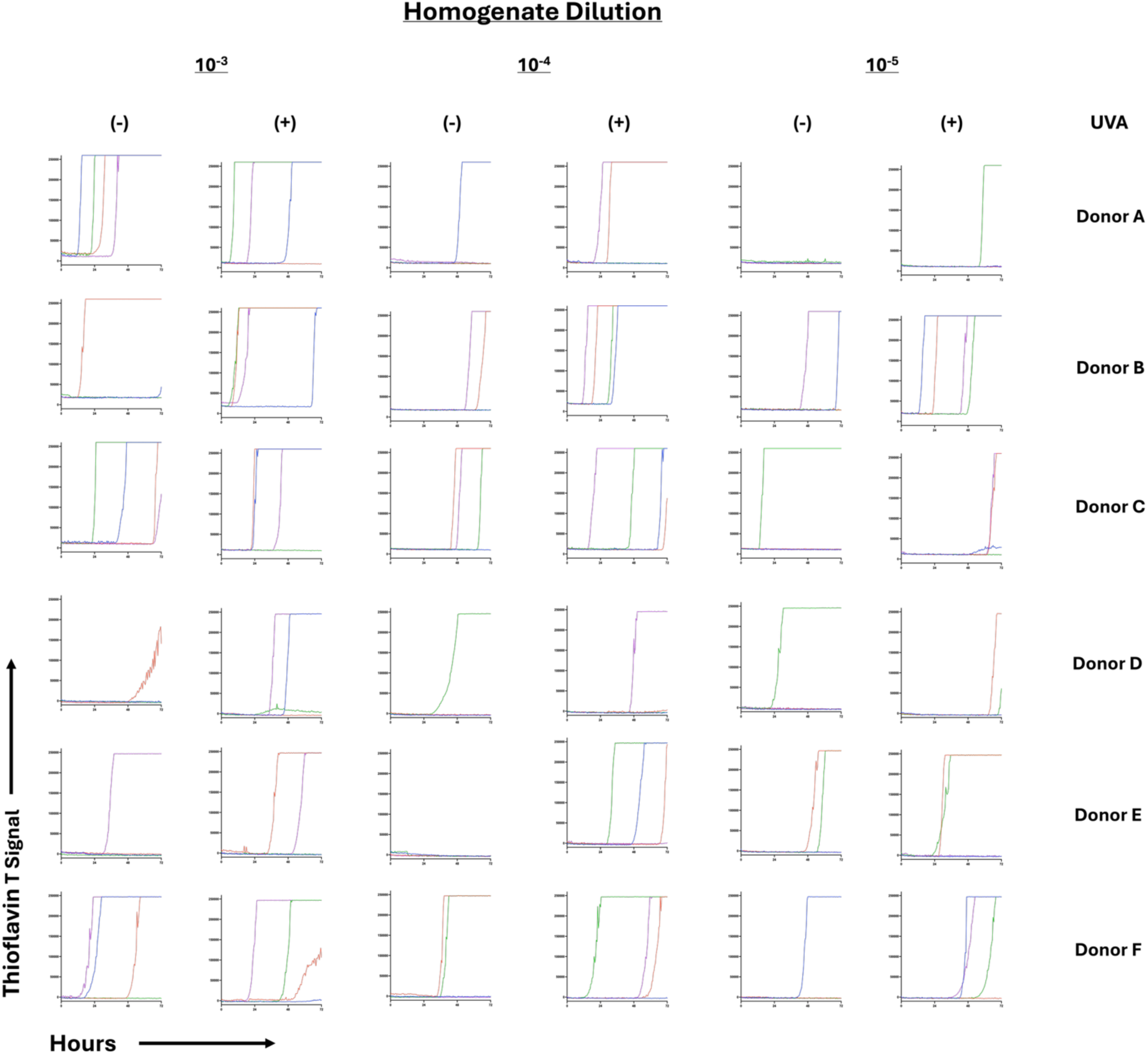
UVA irradiation increases prion-like amyloidogenic activity of cutaneous alpha-synuclein. Discarded abdominal skin from six separate human donors was divided and either irradiated with 5J/cm^2^ UVA daily for 10 days (ex. Peak 368nm) or sham irradiated. Tissue homogenate was then analyzed by the RT-QuIC assay for spontaneous amyloidogenic activity. Thioflavin signal over the 72 hour experiment for each donor +/−UVA is depicted across a 1000-fold dilution series. Each graph represents four technical replicates (identical wells in a 96 well microplate), each depicted in a different color.

**Extended Data Fig. 3:**
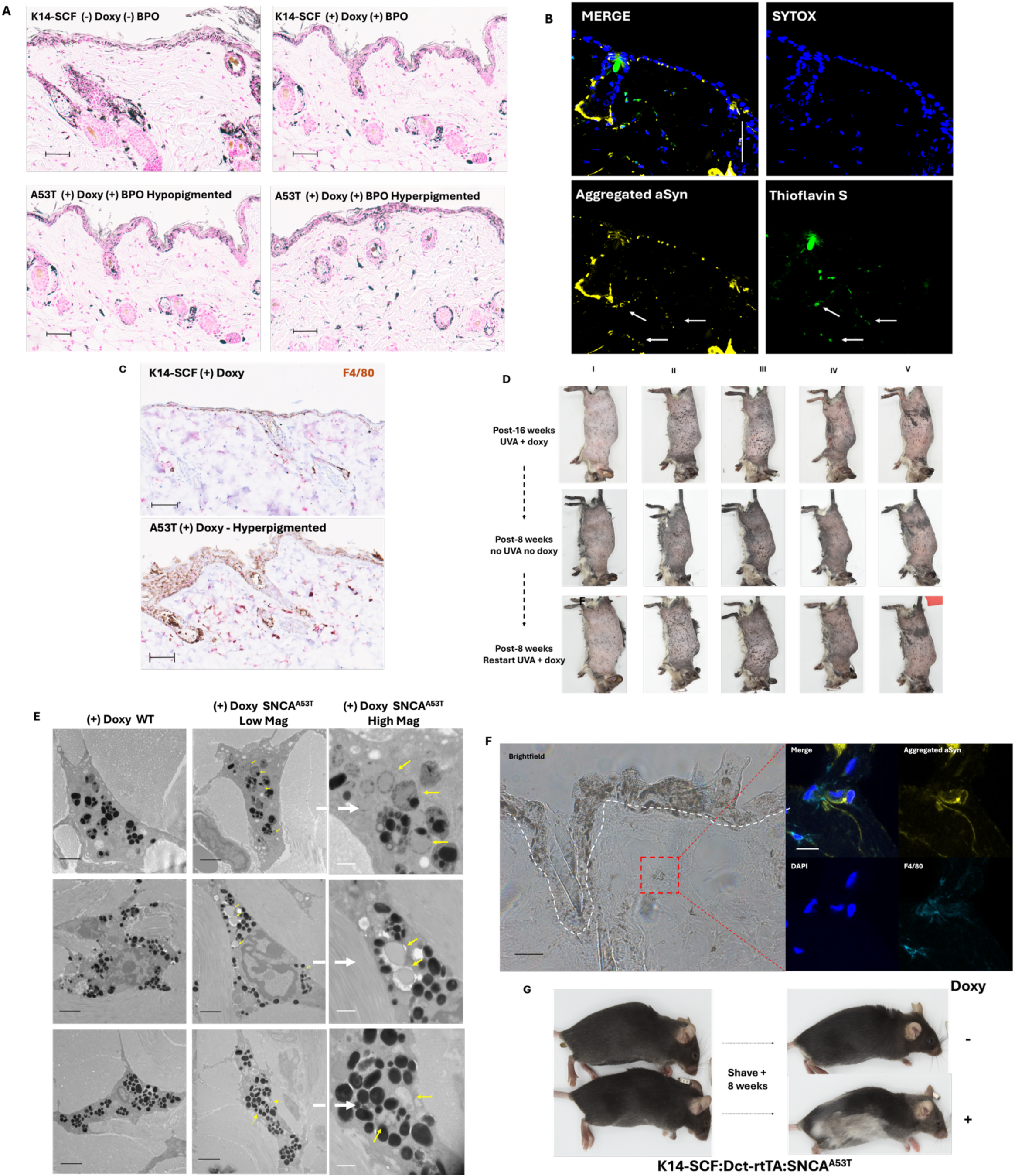
Oxidative stress triggers pigment protein aggregation, resulting in pigmentary dysfunction. (A) Fontana-Masson melanin stain of hyperpigmented SNCA^A53T^ skin shows epidermal and dermal melanin enrichment compared to WT controls. (B) Immunostaining of hyperpigmented mouse skin shows thioflavin-positive dermal aggregated SNCA accumulation. (C) Representative F4/80 immunostaining of hyperpigmented SNCA^A53T^ mouse skin shows significantly more dermal macrophages than WT controls. (D) After spots develop in response to doxycycline induction and oxidative stress, spots gradually fade with cessation of doxycycline and re-darken with reintroduction of doxycycline without additional oxidative stress at the same body sites, n = 5 mice. (E) TEM of dermal melanophages from SNCA^A53T^ mice, identifiable by enclosure of broken down melanosomes within individual phagosomes, shows accumulation of significant electron-lucent material (yellow arrows) not present in WT mouse melanophages analogous to aggregated SNCA immunostain in (F), black scale bar = 2µm, white scale bar = 500nm. (F) Dermal melanophages in SNCA^A53T^ mice also stain positively for aggregated SNCA deposits. (G) SNCA^A53T^ mice (thought not SNCA^WT^ overexpression mice) fed doxycycline chow also show rapid depigmentation of new fur.

**Extended Data Fig. 4:**
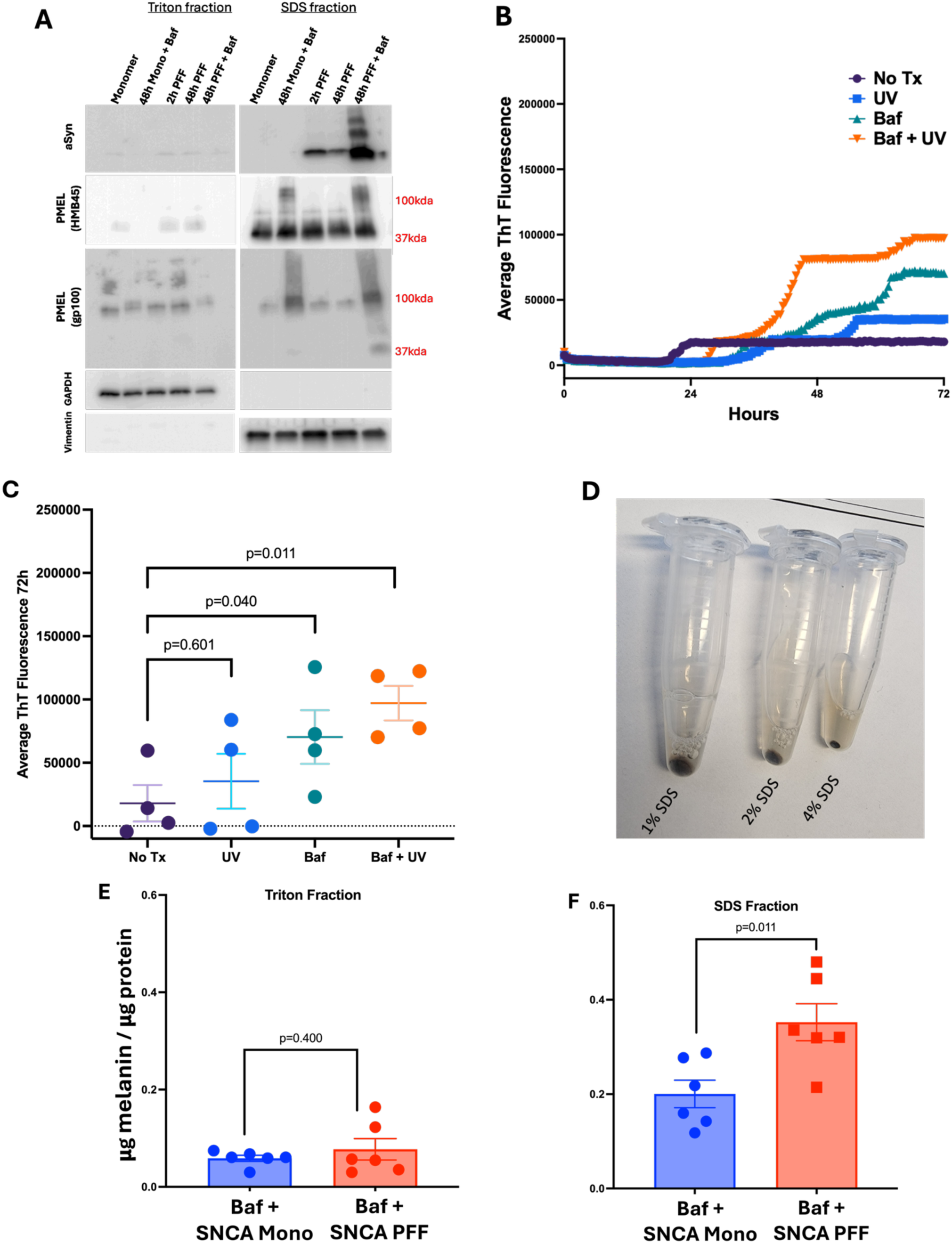
Self-propagating melanosomal protein aggregation contributes to formation of insoluble extracellular melanin deposits. (A) Primary human melanocytes were transfected with 1µg PFFs vs aSyn monomer per 1×10^6^ cells and collected at indicated timepoints. Solubility fractionation showed a significant, time-dependent increase in insoluble SNCA in cells treated with bafilomycin, which was not observed with vehicle control. (B,C) Irradiation of human primary melanocytes with 5J/cm^2^ UVA with or without 100nM bafilomycin led to an increase in aggregation as measured by RT-QuIC primarily in the UV+Baf combination condition (n=4, student’s *t*-test). (D) Solubilization of 1% triton-extracted cell pellets in increasing concentrations of SDS showed progressive pellet dissolution with increasing supernatant melanin turbidity. (E,F) Transfection of primary melanocytes as in (A) with quantification of melanin in the SDS supernatant by absorption at 405nm showed increased partitioning of melanin into the SDS fraction bound to insoluble protein (n=6).

**Extended Data Fig. 5:**
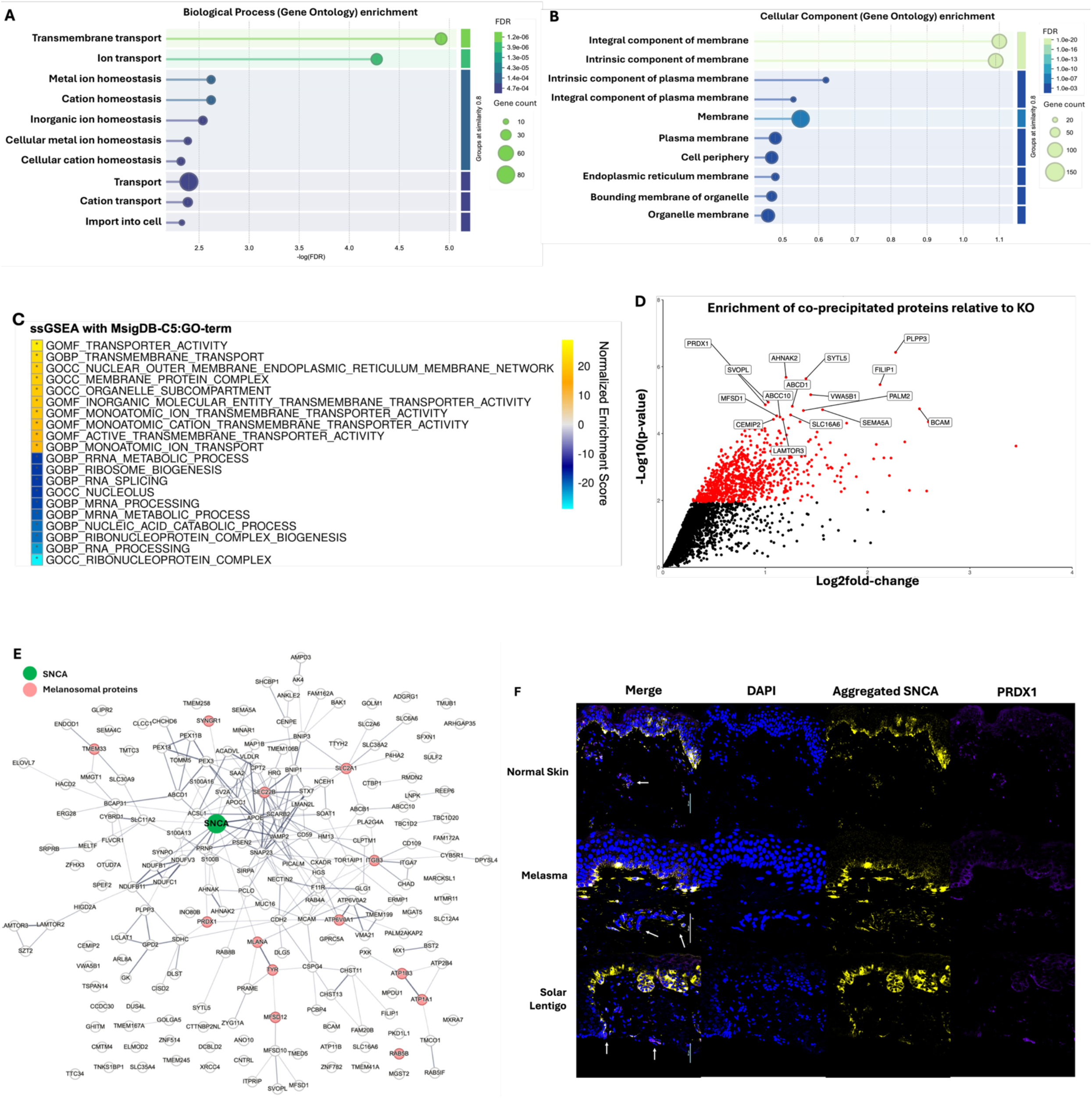
Aggregation partners for alpha synuclein in the melanocytic lineage mirror aggregation partners observed in neurons. (A, B) Gene Ontology (GO) enrichment analysis using the STRING database of proteins with log2(SNCA-overexpressed/SNCA-knockout) > 1 and adjusted p-value < 0.05. Results are visualized for the top 10 terms in the cellular component (A) and biological process (B). (C) Gene Set Enrichment Analysis (GSEA) of the SNCA-interacting proteome using the GO MSigDB C5 gene ontology sets. Results are visualized for the top 20 GO terms. (D) Volcano plot showing the enrichment of protein groups in SNCA-overexpressed compared to SNCA-knockout samples. Labeled proteins represent the top 10 most statistically significant hits based on adjusted p-value. Red dots represent all significant proteins. (E) STRING interaction network of the top 200 most significantly enriched proteins (ranked by adjusted p-value). Each node represents a protein group, and edges indicate known or predicted functional associations. The green node highlights SNCA, and pink nodes denote melanosomal proteins. (F) Immunostain of human hyperpigmentation biopsies showing co-localization of aggregated aSyn with PRDX1 (arrows), which was greatly enriched in the LC-MS screen (white bar = 50µm)

**Extended Data Fig. 6:**
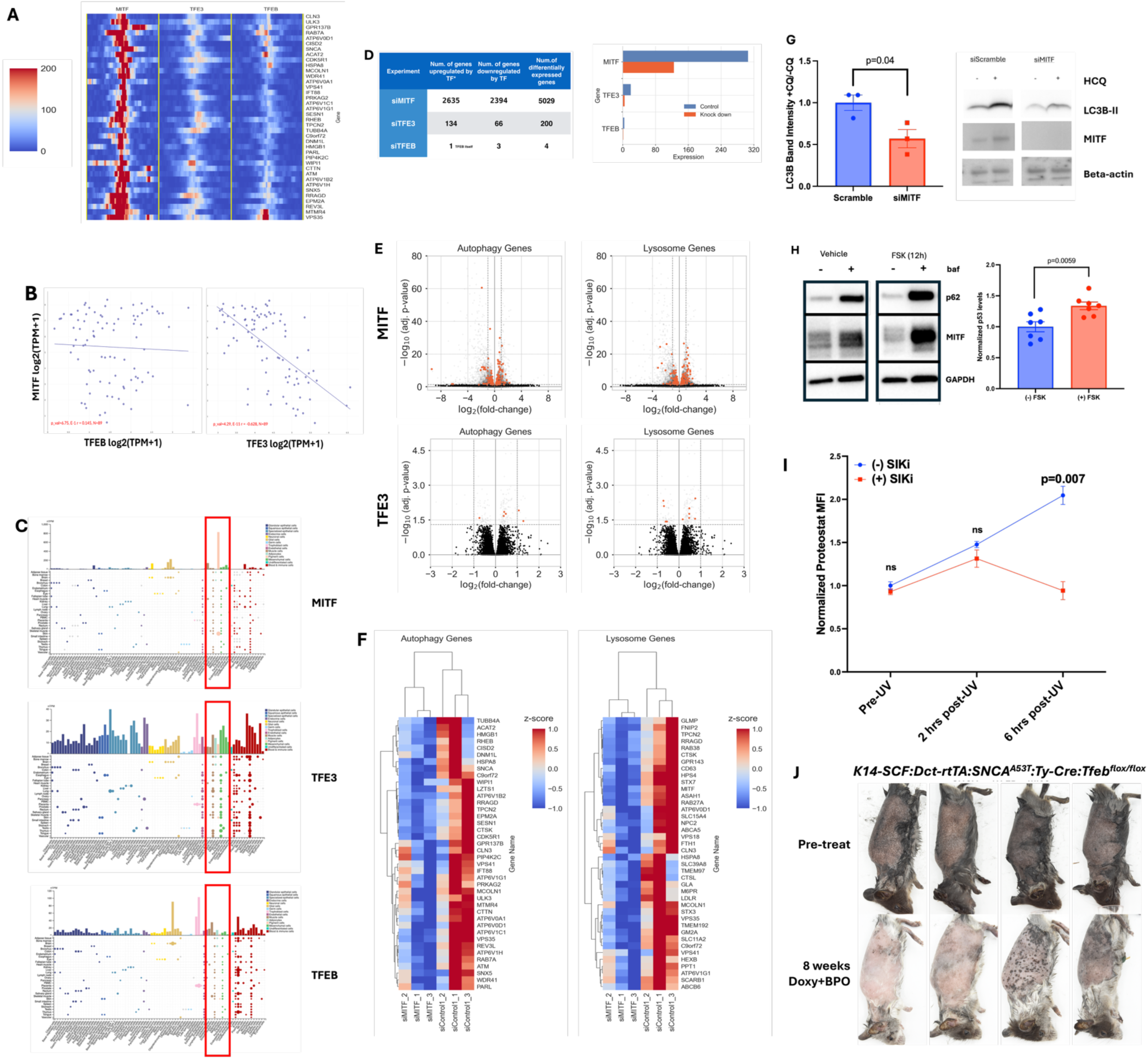
Autophagy is upregulated with the tanning pathway by MITF in the melanocyte lineage. (A) Heatmap plot of MITF, TFE3, and TFEB CUT&RUN signals (normalized by total coverage) in a 4kb region centered at their respective peak summit in select autophagy gene promoters, n = 3 replicates per gene. (B) Transcriptomic data from the CCLE shows no correlation between MITF and TFEB in a cohort of 89 human melanoma tumors but shows significant anticorrelation of MITF and TFE3, which was also observed in our RNA sequencing data. These data also support that TFEB expression is much lower than MITF or TFE3 in the melanocyte lineage. (C) Single-cell RNA sequencing data from The Human Protein Atlas showing that expression of TFE3 is dramatically lower than MITF, and that TFEB expression is minimal in the melanocyte lineage (pink bars in red boxes). (D) Expression levels of MITF, TFE3, and TFEB in control and siMITF, siTFE3, and siTFEB experiments respectively. Samples were cultured in conditions established as activating each respective transcription factor for both target genes and corresponding scramble controls (2 hour nutrient starvation vs. normal media for TFEB and TFE3, and media with or without human melanocyte growth serum for MITF). Expression change between the control and knockdown conditions is statistically significant in all three experiments (p-value = 9.27×10^−8^, 9.26×10^−58^, 8.10×10^−3^, respectively). (E) Volcano plot of differentially expressed autophagy– and lysosome-related genes in siMITF and siTFE3 experiments. (F) Heatmap plot of expression of autophagy-and lysosome-related genes in siMITF experiment. The top 40 differentially expressed genes by adjusted p value relative to scramble control are displayed. (G) Transient siRNA-mediated knockdown of MITF induced a significant decrease in autophagocytic flux in human primary melanocytes as measured by difference in band intensity of LC3B-II with and without 24h 20μM chloroquine, n=3, student’s *t*-test, p<0.05. (H) Activation of the tanning pathway with 20µM FSK increases autophagocytic activity as measured by an increase in p62 levels in forsklolin and bafilomycin-treated cells (n=7, student’s *t*-test). (I) Primary melanocytes cultured in human melanocyte growth supplement-free media for 24 hours with or without SIK inhibitor HG-9-91-01 were irradiated with 5J/cm^2^ UVA (368nm) and misfolded protein was measured by flow cytometry with proteostat dye. SIKi treated cells were able to reduce misfolded proteins levels by 6 hours while starved melanocytes showed continued increase in misfolded protein levels, n=3, student’s *t*-test, p<0.01. (J) 8 week doxycycline chow administration plus 3x/week 10% topical BPO treatment produced similar phenotypes in *Ty-Cre:Tfeb^flox/flox^:K14-SCF:Dct-rtTA:SNCA^A53T^*mice compared to Ty-Cre:TFfeb^WT^*:K14-SCF:Dct-rtTA:SNCA^A53T^*mice, n=4 mice

**Extended Data Fig. 7:**
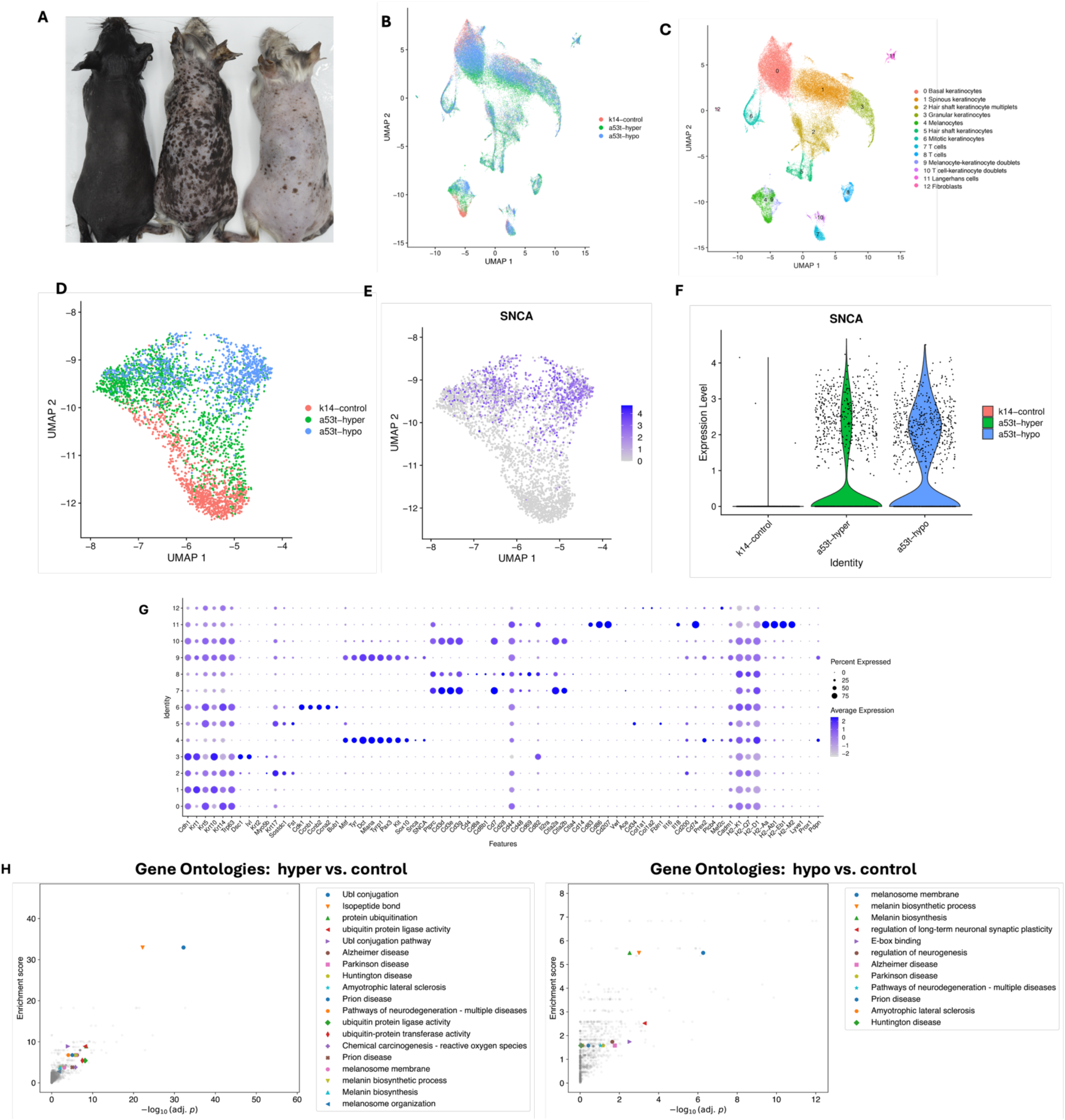
Protein aggregation in melanocytes triggers neurodegeneration-like changes. (A) epidermal cells from 8 month old SNCA^A53T^ homozygous mice and K14-SCF control mice treated for 6 months with 3x/week 10% BPO and induced with doxycycline chow were analyzed. Uniform Manifold Approximation and Projection (UMAP) of all experimental groups (control, hyper, hypo). The cells from the three groups are seen to be well mixed. (C) UMAP of experimental groups (control, hyper, hypo) by cell type. Thirteen clusters were identified using known biomarkers for cell types in the skin (Methods). Only cluster 4, which was identified as melanocytes, was used for downstream analysis. (D-F) Zoomed-in UMAP of (d) experimental groups and (E,F) SNCA expression levels in the melanocyte cluster (cluster 4). Both figures exclude melanocyte-keratinocyte doublets (cluster 9). (G) Dot plot summarizing the cell-type biomarker average log-expression levels and the percentage of expressing cells for individual clusters in the scRNA-seq analysis of human *SNCA*-expressing mice (Methods). (H) Gene Ontology (GO) analysis of differentially expressed genes. GO categories from DAVID are shown as scatter plots for each paired comparison of hyper vs. control (left) and hypo vs. control (right). Each enrichment score corresponds to a functional annotation cluster consisting of multiple related GO terms and pathways, with the individual categories sorted by log-transformed Benjamini-Hochberg adjusted p-value. Salient GO categories are labeled with colored markers. The remaining unlabeled grey background indicates all other terms produced by DAVID.

**Extended Data Fig. 8:**
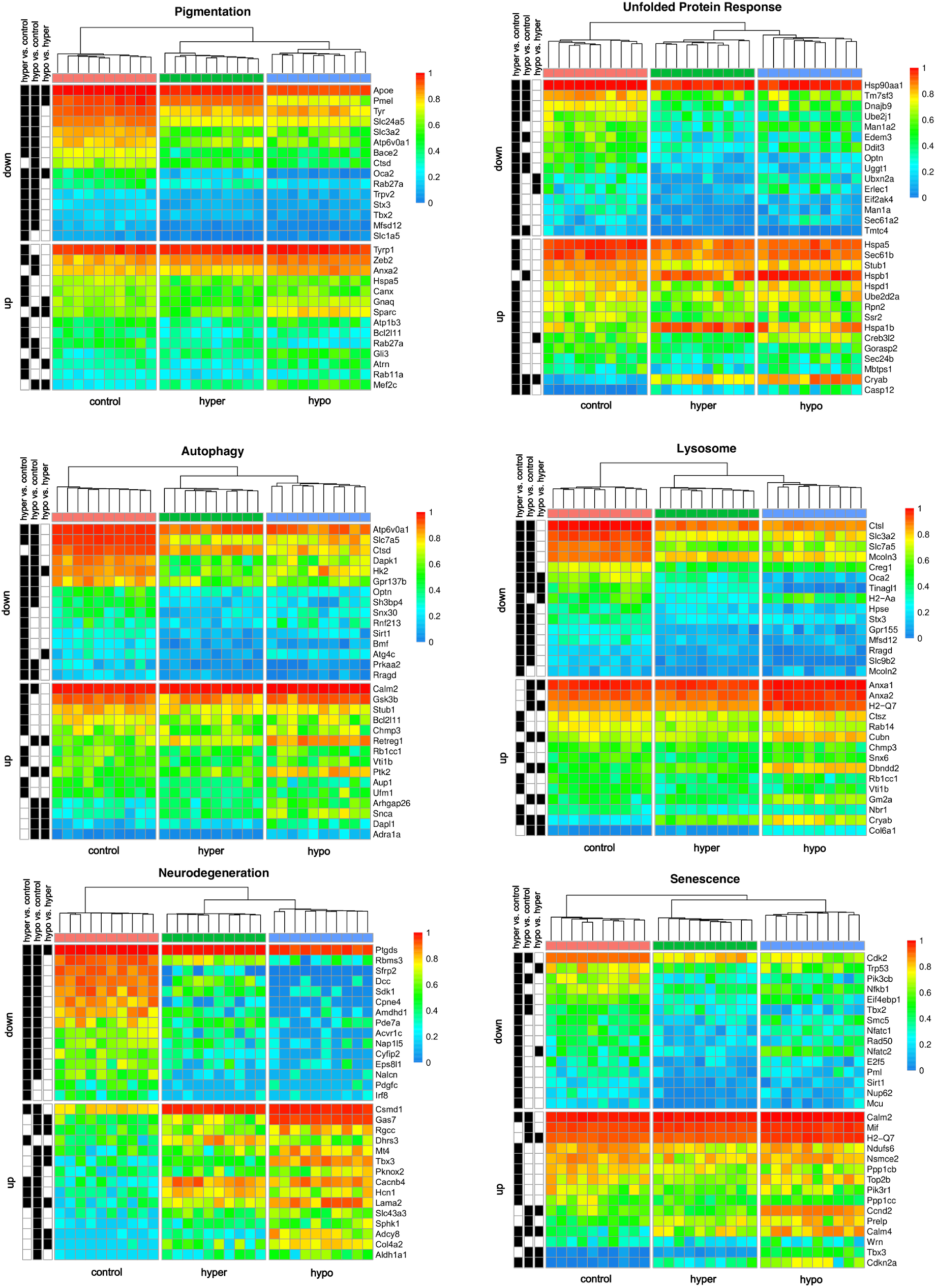
Chronic proteinopathy in melanocytes induces neurodegeneration-like changes and a protective senescent state. Heat maps show unbiased hierarchical clustering of coarse-grained single cells by expression patterns. The single cells were coarse-grained into 10 blocks within each experimental group, and block-averaged expression profiles of GO genes were used to perform hierarchical clustering (Methods). The result correctly reproduced the experimental groups labeled at the bottom and shown by the dendrogram color labels at the top with red, green, and blue corresponding to control, hyper, and hypo, respectively. Black and white row annotations indicate statistical significance and non-significance (Benjamini-Hochberg *p*_*adj*. < 10^−3^, respectively, of differential expression between the indicated groups.

**Extended Data Fig. 9:**
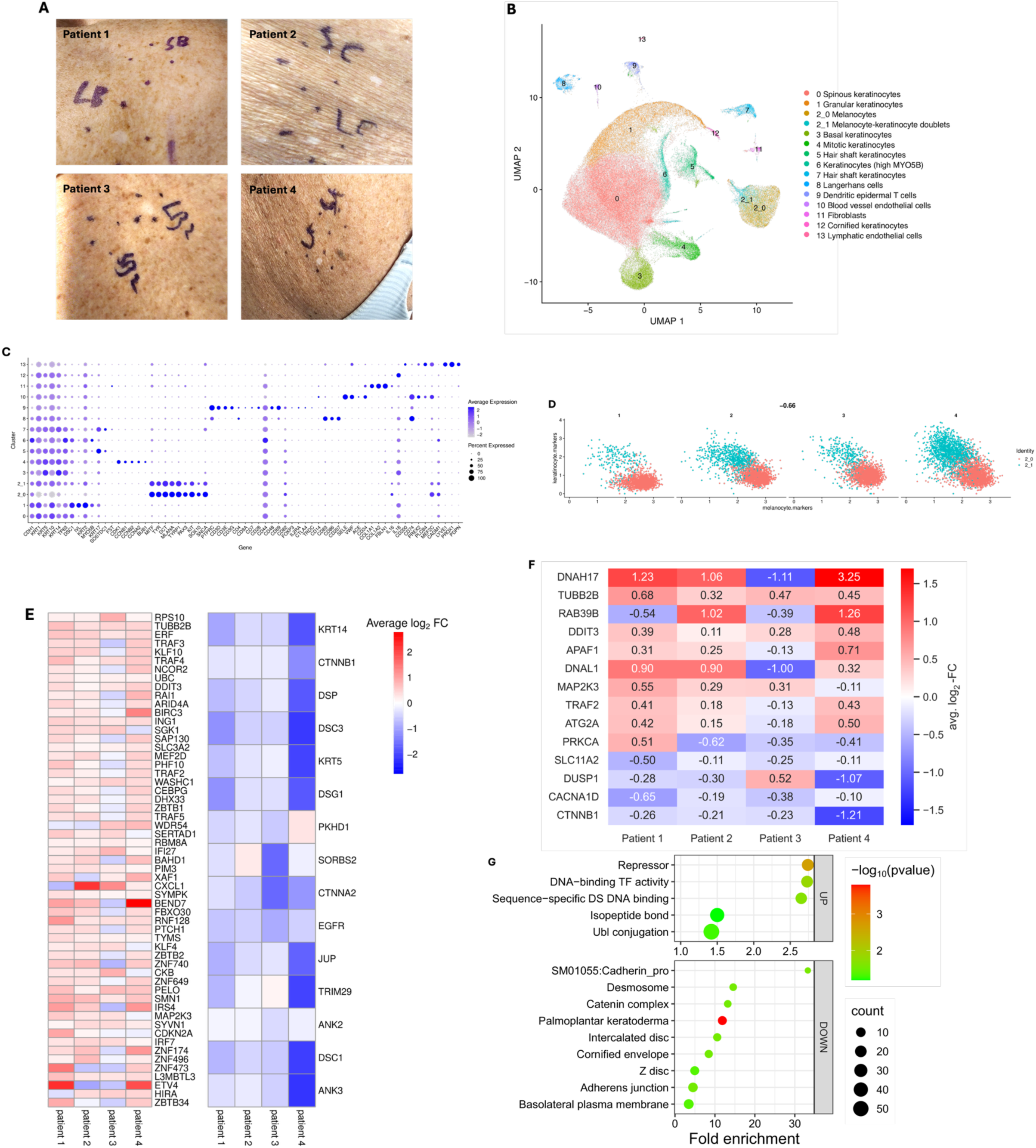
Evidence of chronic proteinopathy and neurodegeneration-like features are seen in photodamaged melanocytes. (A) Representative photographs of IGH lesional vs. perilesional samples and background skin from all four participants. (B) Combined UMAP of single cells from all patients and conditions (lesional and perilesional) and inferred cell-type clusters (Methods). (C) Cell-type biomarker average log-expression levels and percentage of expressing cells for individual clusters. Cluster numbers correspond to those shown in B. Cluster 2 contained melanocytes in the original clustering, and a further subclustering distinguished pure melanocytes (cluster 2_0) from keratinocyte-melanocyte multiplets (cluster 2_1). (Methods). (D) Mean melanocyte biomarker expression versus mean keratinocyte biomarker expression across patients. The scatter plot shows the subclustering of the original melanocyte cluster (2) into pure melanocytes (cluster 2_0) and melanocyte-keratinocyte doublets (cluster 2_1). The x– and y-axes represent the mean expression of melanocyte biomarkers (MITF, TYR, DCT, MLANA, TYRP1, PAX3, KIT, SOX10, SNCA) and keratinocyte biomarkers (CDH1, KRT1, KRT2, KRT5, KRT10, KRT14, KRT17, TP53), respectively. Patients are labeled above each plot. (E) Heatmap of differentially expressed genes in selected GO categories across IGH patients. A subset of the top 300 genes that are up or down regulated in IGH lesional vs.perilesional samples are shown in the left and right heatmaps, respectively. The UP genes (left) include the union of genes from the GO categories associated with CROSSLNK, isopeptide bond, ubl conjugation, znf TRAF, and TNF signaling pathway; the DOWN genes (right) are the union of genes from the GO categories associated with cornified envelope, cell-cell adhesion, desmosome, and structural constituent of cytoskeleton (Supplementary Tables 2 and 3). The colormap indicates average log2 fold change. All reported genes satisfy *padj*. < 10^−3^. (F) Neurodegeneration and PD-related genes show consistent dysregulation in all four IGH patients. Only DE genes (combined adjusted p-value < 10^-3; absolute mean average log_2 FC > 0.2) from the GO categories “Pathways of neurodegeneration – multiple diseases” and “Parkinson disease” are shown. (G) Selected GO terms from gene ontology analysis of the top 300 UP and DOWN gene sets in IGH melanocyte (Methods). Bubble size indicates the number of DE genes associated with the term. The reported GO terms have Benjamini p-value < 0.1.

