## Supplemental Figures for "UV induces common cutaneous amyloid-like melanosomal protein aggregates"

| Category (# Genes) | Gene Sets |
|---|---|
| Pigmentation (191) | pigmentation (GO:0043473), pigment granule (GO:0048770) |
| Neurodegeneration (4003) | Genes4PD |
| Lysosome (595) | lysosome (KEGG:04142), lysosome (GO:0005764) |
| Autophagy (462) | regulation of autophagy (GO:0010506), positive regulation of autophagy (GO:0010508), macroautophagy (GO:0016236), microautophagy (GO:0016237), chaperone-mediated autophagy (GO:0061684), late endosomal microautophagy (GO:0061738), autophagy – animal (KEGG:04137) |
| Senescence (242) | cellular senescence (GO:0090398), stress-induced premature senescence (GO:0090400), oncogene-induced cell senescence (GO:0090402), positive regulation of cellular senescence (GO:2000774), cellular senescence (KEGG:04218) |
| UPR (241) | endoplasmic reticulum unfolded protein response (GO:0030968), response to unfolded protein (GO:006986), cellular response to unfolded protein (GO:0034620), IRE1-mediated unfolded protein response (GO:0036498), protein processing in endoplasmic reticulum (KEGG:04141) |

Supplementary Figure 1: **Curated Gene Ontology Categories for Unbiased Clustering.** The gene sets were categorized according to Gene Ontology (GO) classifications, the Kyoto Encyclopedia of Genes and Genomes (KEGG) pathways, and the Genes4PD study. Parenthetical annotations denote the corresponding GO accession numbers or KEGG pathway identifiers.

| Go Category | Enrichment Score | Adj. $p$-value | Genes |
|---|---|---|---|
| CROSSLNK | 2 | 1.0E0 | ARID4A, BEND7, CEBPG, ETV4, ERF, L3MBTL3, PHF10, RBM8A, SAP130, ING1, NCOR2, PELO, RAI1, SLC3A2, SMN1, SYMPK, TYMS, ZBTB1, ZBTB2, ZBTB34, ZNF174, ZNF473, ZNF496, ZNF649, ZNF740 |
| Isopeptide bond | 2 | 6.3E-2 | ARID4A, BEND7, CEBPG, ETV4, ERF, KLF4, L3MBTL3, PHF10, RBM8A, SAP130, TRAF2, TRAF3, TRAF4, TRAF5, WASHC1, CKB, ING1, IFI27, IRF7, MEF2D, NCOR2, PTCH1, PELO, RAI1, RPS10, SLC3A2, SMN1, SYMPK, TYMS, TUBB2B, UBC, ZBTB1, ZBTB2, ZBTB34, ZNF174, ZNF473, ZNF496, ZNF649, ZNF740 |
| Ubl conjugation | 2 | 5.0E-2 | ARID4A, BEND7, CEBPG, DHX33, DDIT3, ETV4, ERF, FBXO30, KLF10, KLF4, L3MBTL3, PHF10, PIM3, RBM8A, SERTAD1, SAP130, TRAF2, TRAF3, TRAF4, TRAF5, WASHC1, WDR54, BIRC3, BAHD1, CKB, CDKN2A, HIRA, ING1, IRS4, IFI27, IRF7, MEF2D, NCOR2, PTCH1, PELO, RAI1, RPS10, RNF128, SGK1, SLC3A2, SMN1, SYMPK, SYVN1, TYMS, TUBB2B, UBC, ZBTB1, ZBTB2, ZBTB34, ZNF174, ZNF473, ZNF496, ZNF649, ZNF740 |
| Znf TRAF | 1.91 | 2.7E-4 | FBXO30, TRAF2, TRAF3, TRAF4, TRAF5, XAF1 |
| TNF signaling pathway | 1.22 | 4.9E-1 | CXCL1, TRAF2, TRAF3, TRAF5, BIRC3, MAP2K3 |

Supplementary Figure 2: Select GO pathways enriched in the top 300 upregulated genes in IGH lesional vs. perilesional samples sorted by largest $\log_2$ fold change.

| Go Category | Enrichment Score | Adj. $p$-value | Genes |
|---|---|---|---|
| Cornified envelope | 3.25 | 2.9E-2 | DSC1, DSC3, DSG1, DSP, JUP, KRT14 |
| Cell-cell adhesion | 2.78 | 3.2E-1 | PKHD1, CTNNA2, CTNNB1, DSC1, DSC3, DSG1, DSP, EGFR, JUP, TRIM29 |
| Desmosome | 2.78 | 2.9E-2 | DSC1, DSC3, DSG1, DSP, JUP |
| Structural constituent of cytoskeleton | 1.96 | 5.0E-1 | ANK2, ANK3, CTNNA2, DSP, KRT14, KRT5, SORBS2 |

Supplementary Figure 3: Select GO pathways enriched in the top 300 downregulated genes in IGH lesional vs. perilesional samples, sorted by smallest $\log_2$ fold change.
